# Endosomal Sorting Drives the Formation of Mutant Prion Endoggresomes

**DOI:** 10.1101/2020.09.27.315846

**Authors:** Romain Chassefeyre, Tai Chaiamarit, Adriaan Verhelle, Sammy Weiser Novak, Leonardo R. Andrade, Uri Manor, Sandra E. Encalada

## Abstract

Intra-axonal misfolded protein aggregates are a pathological feature of neurodegenerative diseases. How aggregates are formed and cleared is key to maintaining proteostasis. By systematically analyzing the trafficking itinerary of a misfolded GPI-anchored prion protein (PrP) mutant, we unveil endocytic pathways that drive its immediate degradation in the soma, versus its aggregation in axons inside endosomal structures we termed endoggresomes. Axonal sorting occurs post-Golgi, by association of mutant PrP vesicles with Arl8b/kinesin-1/HOPS, a complex that earmarks them for axonal entry, fusion, and aggregation via a mechanism of **<u>a</u>**xonal **<u>r</u>**apid **<u>e</u>**ndosomal **<u>s</u>**orting and **t**ransport-dependent **<u>a</u>**ggregation (ARESTA). Endoggresomes persist in axons due to transport and lysosomal deficits, impairing calcium dynamics and accelerating neuronal death. Reducing ARESTA inhibits endoggresome formation and circumvents these defects. These data identify the endo-lysosomal system as critical for the sorting of misfolded PrP, and ARESTA as an actionable anti-aggregation target that can ameliorate axonal dysfunction in the prionopathies.

## INTRODUCTION

The progressive accumulation of misfolded proteins as intra-neuronal aggregate inclusions is a hallmark of neurodegenerative disorders including Alzheimer’s Disease (AD), Parkinson’s Disease (PD), Huntington’s Disease (HD), and the prion diseases^1^. Axons are particularly vulnerable to the formation of aggregates, which are often observed at sites of axonal swellings, and can be formed from wild-type (WT) or mutant misfolded proteins that lose solubility and become cytotoxic to the neuron and also to neighboring cells via spreading^2^. Despite the importance of aggregates in neurodegeneration, the molecular mechanisms and pathways that tip the balance between protein misfolding versus aggregate clearance resulting in the formation of aggregate inclusions in axons, remain undefined.

To regulate function, polarized neurons rely largely on active microtubule-based transport to localize components to distal axonal domains via kinesin-mediated anterograde movement^3^. As the axon has important but limited lysosomal degradative capacity^4^, long-distance dynein-dependent retrograde transport of axonal cargoes back to the lysosome-enriched soma is critical for their efficient degradation^5, 6^. Neuronal homeostasis also depends on an interconverting endo-lysosomal system that spreads over long axonal distances, to regulate sorting, signaling, and metabolic functions^7^. How dysregulation of these systems results in aggregate formation in axons remains unclear. Disruptions to protein quality control and ubiquitin-proteasome/autophagosome-lysosome clearance pathways can lead to the sequestration of misfolded cytosolic proteins such as tau, *α*-synuclein, TDP-43, FUS, and cytosolic yeast prions, into membrane-free aggresomes, insoluble protein deposits (IPODs), or juxtanuclear quality control (JUNQ) structures^8–14^. Important mechanistic insights on the formation of these cytosolic inclusions have been revealed^9, 15, 16^. However, compelling evidence suggests that accumulation of pathogenic proteins can also occur within the endo-lysosomal system, as several aggregate-prone proteins transit through this system in neurons during processing, and/or as they accumulate within vesicular compartments^17^. How endomembrane proteins such as glycosylphosphatidylinositol (GPI)-anchored proteins form intra-neuronal aggregate inclusions, remains virtually unexplored.

The cellular prion protein (PrP^C^-cellular or PrP^WT^) is a secreted GPI-anchored protein that when misfolded acquires a partially proteinase K-resistant conformation called PrP scrapie (PrP^Sc^), which is associated with toxic pathologies in prion disorders^18–24^. While mostly sporadic, some prion diseases are transmissible, and approximately 10-15% are caused by mutations in the PrP gene, *PRNP* ^25, 26^. Intraneuronal aggregate inclusions have been observed in patient brains with familial PrP mutations, and intra-axonal aggregates have been observed in cellular and mouse familial models of prion disease, including one harboring a nine octapeptide repeat insertion called PrP^PG14^ shown to associate with dementia and ataxia^26–36^. Where and how PrP aggregates form within neurons, remains unclear. PrP^Sc^ has been reported to form aggresomes following proteasome inhibition and after retrotranslocation to the cytosol^37^. However, misfolded PrP has also been observed to accumulate within membrane compartments, pointing toward a role for endosomal trafficking in its aggregation^38^. Indeed, the membrane association and secretion of PrP via its GPI anchor and signal peptide indicate that sorting and trafficking are central to PrP physiology and pathology^39^. In its native or non-native forms PrP traffics to the cell surface from the ER via the Golgi in the lumen of membrane compartments, from where it is endocytosed back for recycling in endosomes, or transits to lysosomes for degradation^18, 39–46^. In non-neuronal cells, this pathway called rapid ER stress-induced export (RESET), was shown to activate within minutes of ER stress to ensure the rapid lysosomal clearance of virtually all misfolded mutant PrP particles^47, 48^. It is unknown whether neurons utilize RESET or other mechanisms to clear misfolded PrP, but its accumulation in axons^49^ suggests that it can escape clearance in the soma. Consistent with this possibility, our earlier work shows the active transport of post-Golgi PrP^WT^ vesicles in and out of the axon by kinesin and dynein^50^, where it may undergo further sorting.

The trafficking and sorting of endo-lysosomes is orchestrated largely by Arl8b, an Arf-like (Arl) family small GTPase that recruits kinesin, and the multi-unit homotypic fusion and protein sorting (HOPS) complex onto late endosomes (LEs), to drive their motility to the cell periphery and towards homotypic fusion with other LEs^51–55^. Arl8b depletion leads to perinuclear lysosome clustering that has been associated with reduced levels of pathogenic proteins in non-neuronal cells^56, 57^, but whether and how Arl8b could regulate aggregate formation in axons is unclear. In this study, by systematically analyzing the trafficking itineraries of the misfolded PrP^PG14^ mutant in mammalian neurons, we unveil endo-lysosomal pathways that impose dramatically different fates on mutant PrP: one drives it for immediate degradation within lysosomes in the soma, while the other shuttles mutant PrP into the axon for aggregation in endoggresomes, a new type of aggregate structure that forms inside endosomal compartments and is associated with neuronal toxicity. Endoggresome formation in axons occurs as a result of the association of Arl8b/kinesin-1/HOPS with Golgi-derived mutant PrP vesicles, sorting them for axonal entry, homotypic fusion, and aggregation via a mechanism we call **<u>a</u>**xonal **<u>r</u>**apid **<u>e</u>**ndosomal **<u>s</u>**orting and **t**ransport-dependent **<u>a</u>**ggregation (ARESTA). Modulating ARESTA inhibits endoggresome formation and prevents neuronal dysfunction, providing an anti-aggregation target that can modulate neuronal dysfunction in the prionopathies.

## RESULTS

### Stress-induced lysosomal clearance of a prion protein mutant in mammalian neurons

Formation of intracellular inclusions could result from an imbalance of synthesis versus clearance pathways. In non-neuronal cells, misfolded GPI-anchored PrP undergoes clearance in lysosomes following cell surface access within minutes of ER stress via RESET^47^. To test whether misfolded PrP^PG14^ is degraded in neurons following this itinerary, we established an *in cellulo* system in dissociated primary murine hippocampal neuronal cultures by transiently expressing fluorescently (mCherry [mCh], EGFP, or mTagBFP2)-tagged or untagged PrP^PG14^ fusions (or PrP^WT^ as control; **Fig. 1a**), shown previously to be properly processed *in vivo*^27, 36, 58–60^. Expression of PrP was under the MoPrP.Xho promoter^61^, and contained the endogenous PrP secretory signal sequence (SS) and GPI anchor sequence to ensure proper translocation to the ER and delivery to the plasma membrane, respectively^62^. Consistent with earlier reports^36, 63^, PrP^PG14^-mCh expressed in Neuro-2a (N2a) cells exhibited mild resistance to proteinase K (PK) as compared to PrP^WT^-mCh, suggesting that this mutant partly and constitutively misfolded and misassembled upon transient expression (**Supplementary Fig. 1a**).

**Figure 1.**
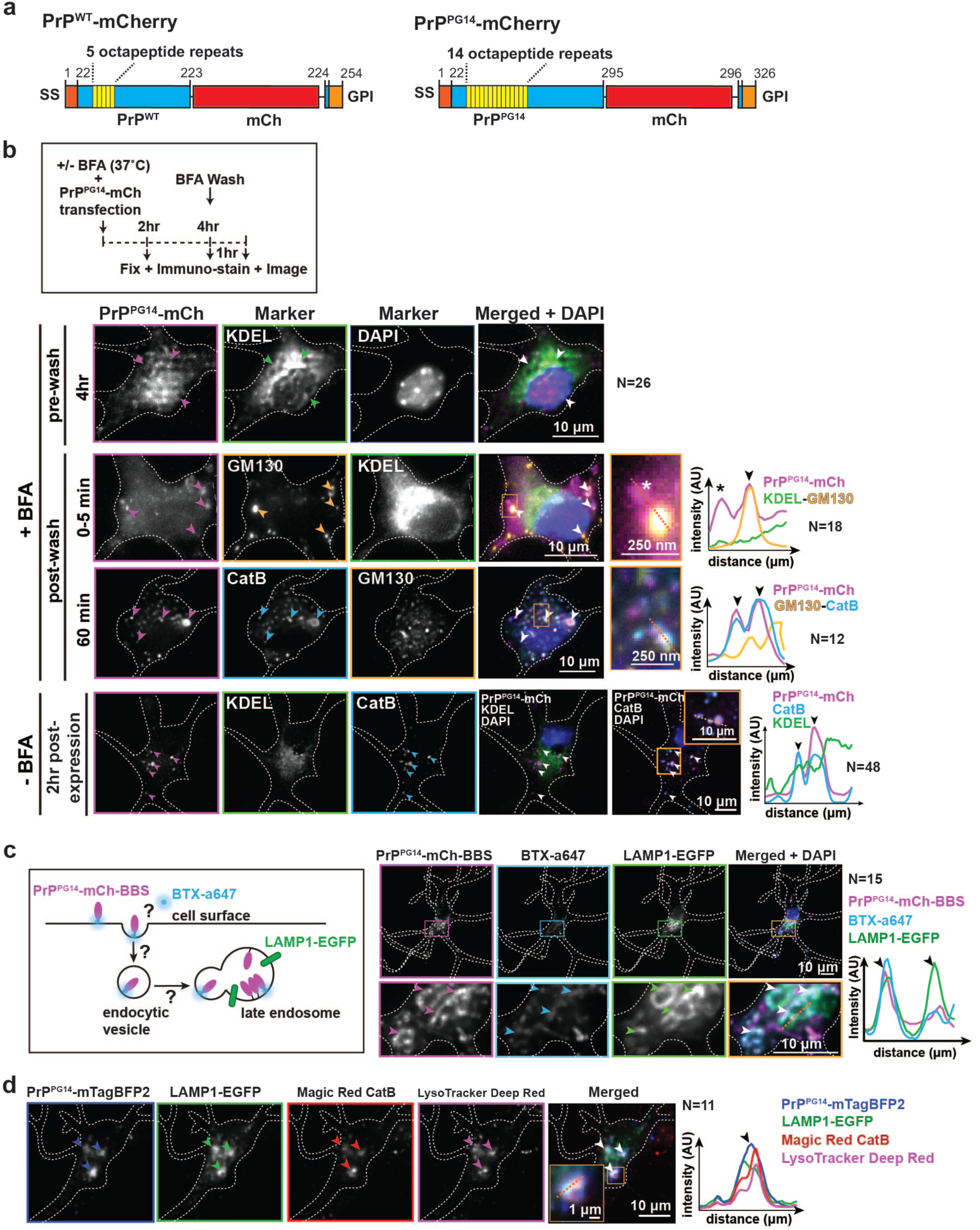
**Neuronal PrP^PG14^ undergoes lysosomal clearance. a**, Schematic of PrP^WT^-mCh and PrP^PG14^-mCh constructs. SS: signal sequence, GPI: GPI anchor. **b**, Experimental outline of BFA assay (top). Representative images of soma of different neurons expressing PrP^PG14^-mCh and stained with antibodies against ER (KDEL), Golgi (GM130), lysosomes (CatB) or with DAPI nuclear marker at indicated time points and conditions (bottom). **c**, Outline of PrP^PG14^-mCh-BBS internalization assay (left). Representative images of the soma of a neuron cotransfected with PrP^PG14^-mCh-BBS and LAMP1-EGFP, and labeled with BTX-a647 (right). **d**, Representative images of the soma of a neuron cotransfected with PrP^PG14^-mTagBFP2 and LAMP1-EGFP, and treated with Magic Red and LysoTracker Deep Red). Arrowheads and asterisk point to colocalization or non-colocalization events, respectively, also shown in enlarged insets and line scan intensity profiles. See also **Supplementary Figure 1**.

To determine whether PrP^PG14^ follows RESET for clearance in neurons, we tested for four characteristic features upon PrP^PG14^-mCh expression: (1) increased ER stress; (2) ER-to-Golgi PrP^PG14^-mCh release; (3) transient PrP^PG14^-mCh access to the cell surface, followed by (4) PrP^PG14^-mCh targeting to lysosomes for degradation. First, co-expression of PrP^PG14^-EGFP and an ER stress transcriptional reporter (ERSE-mCh)^64^ resulted in significantly increased signal compared to that observed in neurons expressing either soluble EGFP or PrP^WT^-EGFP (**Supplementary Fig. 1b**), indicating constitutive ER stress activation. Next, we tested for ER-to-Golgi transit by imaging neurons treated or not with brefeldin A (BFA), an inhibitor of the secretory pathway^65^. Following BFA treatment, PrP^PG14^-mCh was retained in the ER for at least 6 hours (**Fig. 1b and Supplementary Fig. 1c**). However, imaging of untreated neurons or of those fixed at various time points post-BFA wash showed rapid, efficient, and successive translocation of misfolded PrP^PG14^-mCh from the ER to the Golgi, and to compartments positive for Cathepsin B (Cat B), an acidic hydrolase required for lysosomal substrate degradation (**Fig. 1b and Supplementary Fig. 1c**). PrP^PG14^-mCh was highly colocalized with puncta positive for Cat B within 2 hours of its expression in untreated neurons (**Fig. 1b**), indicating that ER export occurred within an acute time period shortly after PrP^PG14^-mCh expression, consistent with non-neuronal RESET dynamics^47^.

To next test whether misfolded PrP^PG14^ transiently accessed the cell surface prior to internalization into LEs, neurons were transfected with PrP^WT^- or PrP^PG14^-mCh fusion constructs containing a Bungarotoxin Binding Sequence (BBS; referred to as PrP^WT^- or PrP^PG14^-mCh-BBS; **Supplementary Fig. 1d**), and treated in the media with AlexaFluor 647-labeled bungarotoxin (BTX) (BTX-a647; **Fig. 1c**)^66^. Live imaging showed colocalization of PrP^PG14^-mCh-BBS puncta with BTX-a647 and with LAMP1-EGFP, a LE/lysosomal resident transmembrane protein (**Fig. 1c**), indicating internalization of cell-surface labeled PrP^PG14^-mCh-BBS into LEs. This labeling was specific, as neurons transfected with PrP^PG14^-fusion constructs without the BBS tag did not reveal any BTX-a647 signal (**Supplementary Fig. 1e**). Presence of PrP^PG14^-mCh at the cell surface was confirmed by immunofluorescence of fixed unpermeabilized neurons with an antibody against mCh (**Supplementary Fig. 1f**). Consistent with an earlier report^67^, PrP^PG14^-mCh membrane signal was weaker than that of PrP^WT^-mCh, suggesting the limited delivery of this mutant to the cell surface, or a short residence time there prior to internalization^48^.

Colocalization of PrP^PG14^-mCh with Cat B indicates its residence in degradative compartments. To further test whether PrP^PG14^ was actively degraded, we imaged neurons expressing PrP^PG14^-mTagBFP2 and LAMP1-EGFP with various acidic and degradative organelle markers. We observed colocalization of PrP^PG14^-mTagBFP2 signal with the acidic-organelle dye LysoTracker and with Magic Red, a fluorogenic substrate that activates upon cleavage by active Cat B^68^ (**Fig. 1d**). Furthermore, live-imaging showed the active disappearance of PrP^PG14^-mCh puncta that colocalized with LAMP1-EGFP, indicating active degradation (**Supplementary Fig. 1g and Video 1**). Altogether, these experiments demonstrate that following ER-stress and a transient excursion to the cell surface, ER export is a pathway involved in the secretion and clearance of misfolded PrP^PG14^ in lysosomes in the neuronal soma, similar to RESET in non-neuronal cells^47^.

### Misfolded PrP^PG14^ forms persistent intra-axonal aggregate inclusions

Having shown that misfolded PrP^PG14^ can be cleared from the neuronal soma (**Fig. 1**), it was unclear how intra-axonal inclusions could form and/or persist in axons, as previously reported^36^. To investigate the fate of axonal misfolded PrP^PG14^, we first determined the localization of PrP^PG14^ in axons. Consistent with previous reports showing PrP^PG14^ limited localization at the cell surface^67^, and with observations in the soma (**Fig. 1c**), immunofluorescence analysis with or without permeabilization using an anti-PrP antibody (D13) showed scarce PrP^PG14^-mCh signal at the axonal cell surface compared to PrP^WT^-mCh (**Fig. 2a**), suggesting that misfolded PrP^PG14^-mCh was either not efficiently delivered to the axonal membrane, or that it is only transiently localized there. We next tested whether intra-axonal inclusions formed in axons of cultured primary neurons transiently expressing PrP^PG14^. Live imaging showed prominent inclusions (defined in **Methods**), one day after PrP^PG14^-mCh transfection, and these increased over time (**Fig. 2b,c**). In contrast, inclusions were largely absent in axons of neurons expressing PrP^WT^-mCh. Co-expression drove the accumulation of PrP^WT^-mCh with PrP^PG14^-EGFP-inclusions, whereas no accumulations were observed in axons of neurons co-transfected with PrP^WT^-EGFP and PrP^WT^-mCh (**Supplementary Fig. 2a**). These observations suggest that PrP^PG14^-EGFP promotes the coalescence of PrP^WT^- mCh, and/or that misfolded PrP^PG14^-EGFP molecules might act as seeds to induce a conformational conversion of PrP^WT^-mCh to promote its aggregation. Together with observations showing partial PrP^PG14^-mCh PK resistance (**Supplementary Fig. 1a**), these data suggest that misfolded PrP^PG14^ forms aggregates in axons, thus we henceforth refer to these as aggregate inclusions or aggregates.

**Figure 2.**
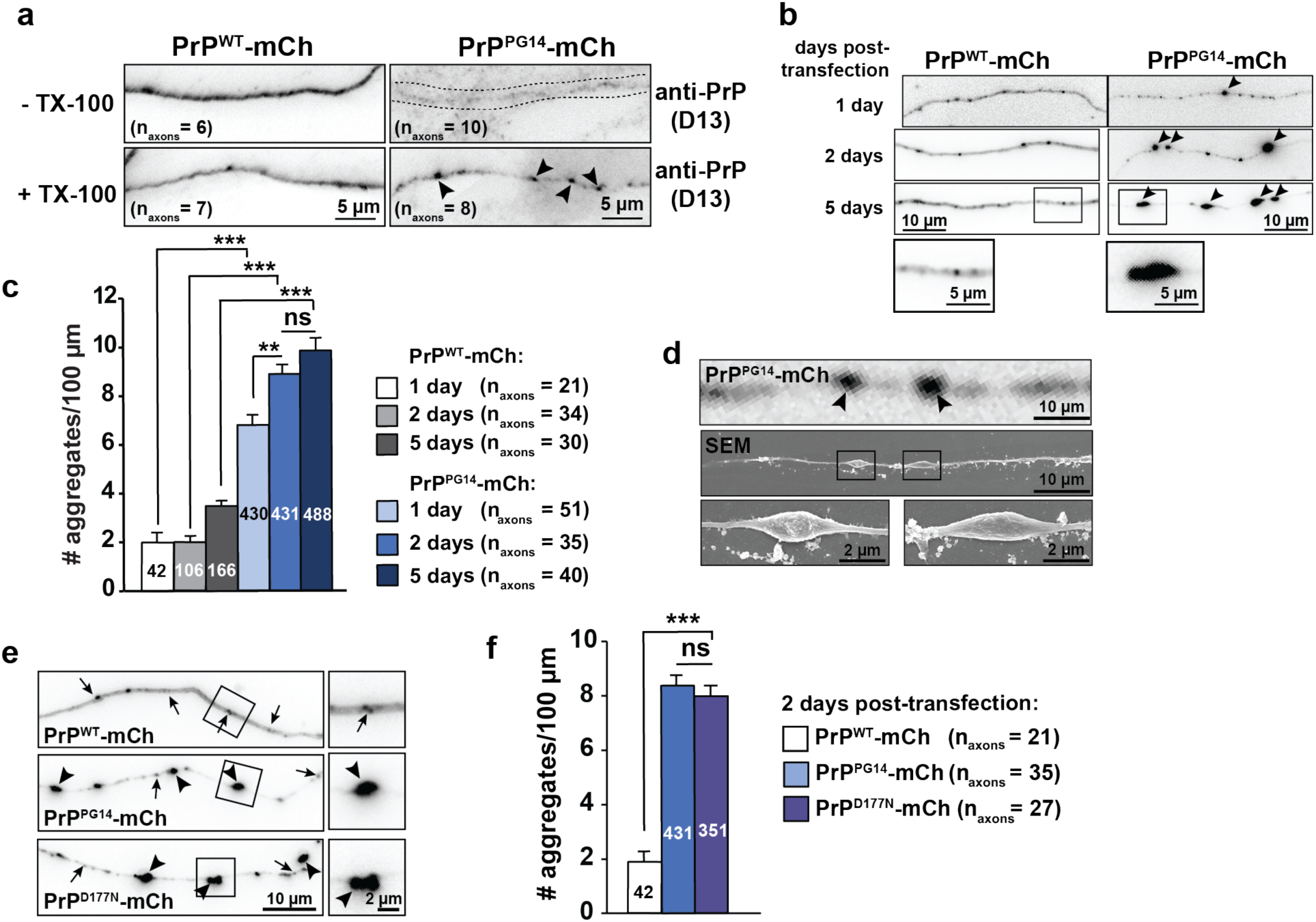
**PrP^PG14^ forms intra-axonal aggregates and swellings**. **a**, Immunofluorescence images of axons expressing PrP^WT^-mCh and PrP^PG14^-mCh in the presence or absence of detergent (TX-100). Arrowheads point to aggregates. **b**, Representative images of axons expressing PrP^WT^-mCh and PrP^PG14^-mCh. Arrowheads point to aggregates. Insets show enlargements. **c**, Quantitation of aggregate densities from (**b**). N_aggregates_ shown inside bars. **d**, Correlative fluorescence (top) and SEM (middle-bottom) images of PrP^PG14^-mCh aggregate swellings. N_swellings_=20. Arrowheads point to aggregates. Insets show enlargements. **e**, Representative images of axons expressing PrP^WT^-mCh, PrP^PG14^-mCh, or PrP^D177N^-mCh 2 days post-transfection. Arrows point to vesicles. Arrowheads point to aggregates. **f**, Quantitation of aggregate densities from (**e**). N_aggregates_ shown inside bars. All values are shown as mean ± SEM. **p<0.01, ***p<0.001, ns = not significant, Student’s t-test, Šidák correction (**c,f**). See also **Supplementary Figure 2.**

To further characterize mutant PrP aggregates, we did correlative fluorescence and scanning electron microscopy (SEM), and observed that they localized to axonal dystrophic swellings averaging 1.57 ± 0.13 µm and 0.54 ± 0.05 µm in length and width, respectively (**Fig. 2d and Supplementary Fig. 2b,c**). Aggregates also formed to the same degree in axons of neurons expressing untagged PrP^PG14^ or PrP^PG14^-EGFP (**Supplementary Fig. 2d,e**), in differentiated N2a cells (**Supplementary Fig. 2f**), and in neurons expressing PrP^D177N(M128)^-mCh (**Fig. 2e,f**), a point mutation (D178N/M129 in humans) that causes fatal familial insomnia, a human prion disease^69^. Altogether, these results indicate that while PrP^PG14^ is degraded in the soma (**Fig. 1**), it forms aggregates at axonal swellings sites that persist over time, suggesting the presence of mechanisms that actively promote aggregate formation and/or of those that impair their clearance.

### Post-Golgi PrP^PG14^ vesicles and aggregates are transported and accumulate within endo-lysosomal compartments in axons

In addition to large aggregates, live imaging using pseudo-total internal reflection fluorescence microscopy (TIRFM) revealed the active bidirectional transport of PrP^PG14^- mCh vesicles in axons (**Fig. 3a and Videos 2,3;** vesicles versus aggregates defined in **Methods**). These observations suggested the possibility that PrP^PG14^-mCh vesicles transiting within the secretory/endosomal system contributed to the formation of larger aggregates. To test this, we used high-resolution single-particle image analysis^70^ to first characterize the dynamics of PrP^PG14^ vesicle transport in axons. The majority of motile PrP^PG14^-mCh vesicles exhibited an anterograde movement bias similar to PrP^WT^-mCh vesicles (**Supplementary Fig.3a and Movie 2**), and of previously characterized YFP-PrP^C^ vesicles^50^, mobilizing towards the synapse with fast velocities consistent with kinesin-mediated transport (**Fig. 3a,b and Supplementary Fig. 3b**). In contrast, all PrP^PG14^-mCh aggregates remained largely stationary, or moved with net velocities that were an order of magnitude (20-50 times) slower (0.01-0.5 µm/sec) than those of vesicles (1-4 µm/sec) (**Fig. 3a,b and Supplementary Fig. 3b**), suggesting that aggregates and vesicles are two differentially transported cargo populations.

**Figure 3.**
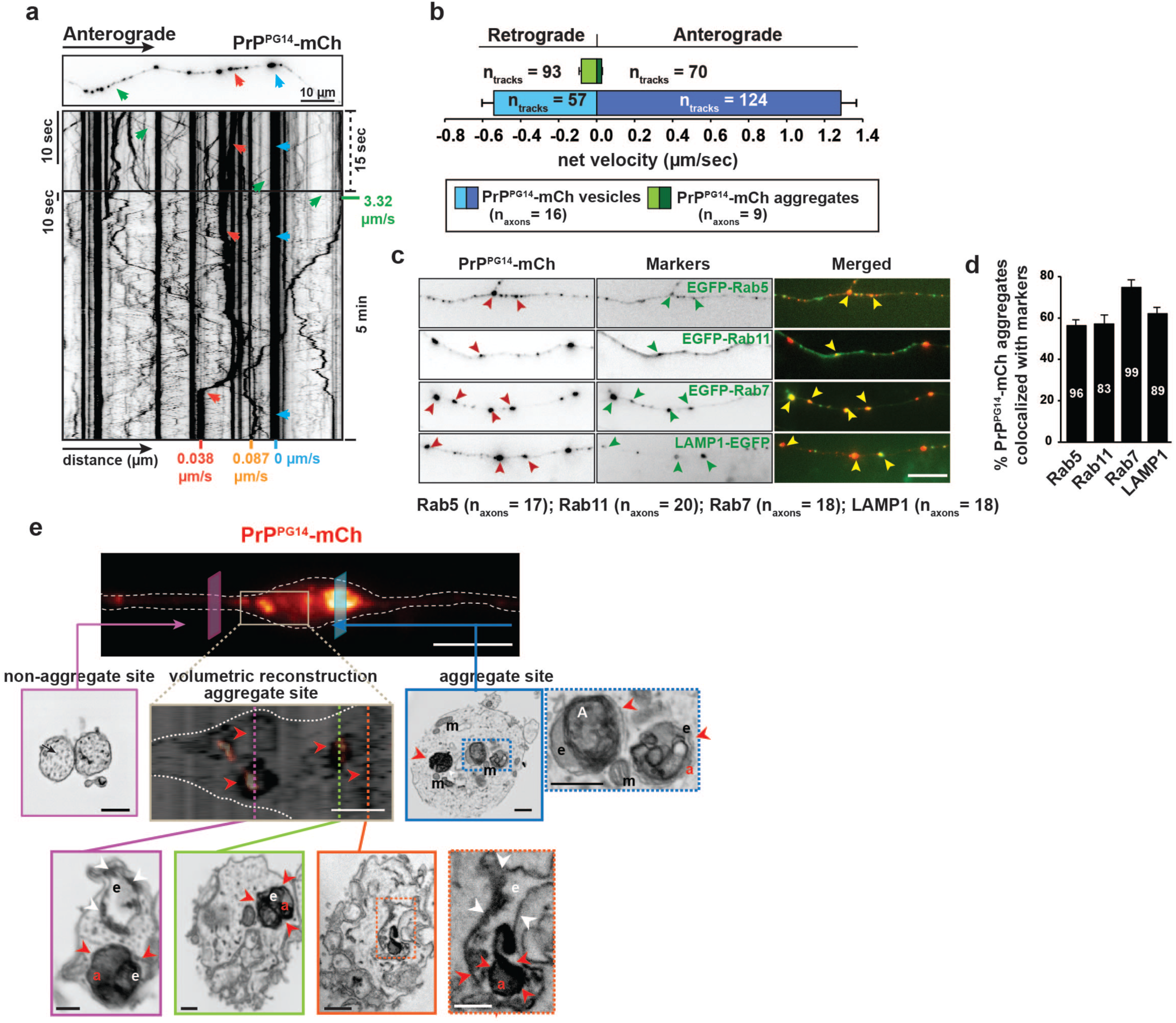
**PrP^PG14^ is transported and accumulates within endo-lysosomal compartments in axons. a**, First frame of movie of an axon expressing PrP^PG14^-mCh (top), and corresponding kymographs (middle-bottom). Arrows point to vesicles or aggregates moving in the anterograde (green) or retrograde (red) directions, or that are stationary (blue). **b**, Average net velocities of PrP^PG14^-mCh vesicles and aggregates. **c**, Representative images of axons co-expressing PrP^PG14^-mCh and markers for early endosomes (EGFP-Rab5), recycling endosomes (EGFP-Rab11), LEs (EGFP-Rab7), and endo-lysosomes (LAMP1-EGFP). Arrowheads point to colocalizing aggregates. Scale bar = 10 μm. **d**, Quantitation of aggregate densities from (**c**). N_aggregates_ shown inside bars. All values are shown as mean ± SEM. **e**, Correlative fluorescence (top) and Serial Sectioning Scanning Electron Microscopy (S3EM) representative cross sections (bottom). Volumetric 3D reconstruction through axonal PrP^PG14^-mCh aggregate site is shown with super-resolution radial fluctuation (SRRF) fluorescence overlay. Red arrowheads point to PrP^PG14^-mCh-positive aggregates within endosomal membranes. White arrowheads point to intraluminal vesicles. m = mitochondria; e = endosome; a = aggregate. Scale bar of top panel of correlative image = 10 μm. All other scale bars = 500 nm. See also **Supplementary Figure 3.**

To identify the type of compartment containing PrP^PG14^, we tracked the trajectories of PrP^PG14^ vesicles and aggregates together with those of various secretory and endo-lysosomal cargoes. Live time-lapse showed significant cotransport of PrP^PG14^ fast-moving vesicles with a Golgi-derived vesicle marker EGFP-Neuropeptide Y (NPY)^71^, and of vesicles and larger aggregates with EGFP-labeled Rab5, Rab11, Rab7, and LAMP1, markers for early, recycling, LEs, and endo-lysosomes, respectively (**Fig. 3c,d and Supplementary Fig. 3c**). Both vesicles as well as large aggregates cotransported and/or colocalized with endo-lysosomal markers, consistent with the notion that PrP^PG14^ traffics and accumulates within endomembranes. To further probe this possibility, we imaged the live time-lapse transport of a photoactivatable (PA) PrP^PG14^-mCh fusion following irradiation of the neuronal soma with a 405 nm laser, in neurons also expressing EGFP-Rab7. We observed the rapid transition of PrP^PG14^-PAmCh from the Golgi to EGFP-Rab7-positive LEs, including to large aggregate structures, within 60 minutes post-photoactivation (**Supplementary Fig. 3d**). To directly test whether PrP^PG14^ aggregates localized within endomembranes we performed correlative light microscopy and serial or single cross-section scanning electron microscopy (S3EM) of aggregate versus non-aggregate regions. Volumetric 3D reconstructions of serial sections of aggregate sites allowed the accurate mapping of PrP^PG14^-mCh signal to structures in the S3EM images (**Supplementary Fig. 3e**). Compared to non-aggregate regions, cross-sections that mapped to PrP^PG14^-mCh-positive aggregate swelling sites revealed large electron-dense structures that resided within membrane compartments resembling endosomes (**Fig. 3e**). Collectively, these observations indicate that PrP^PG14^ is actively transported in Golgi-derived endosomal compartments. In the axon, PrP^PG14^ can accumulate inside larger aggregate endomembrane structures that we name the endoggresome.

### Kinesin-1-mediated anterograde transport of PrP^PG14^ endo-lysosomal compartments drives endoggresome formation in axons

To probe into the origin of endoggresomes, and to test whether PrP^PG14^ vesicles contribute to their formation in axons, we compared the transport dynamics of PrP^PG14^-EGFP vesicles and aggregates in wild-type (WT) axons and those from kinesin-1C (KIF5C) knockout (*Kif5c^-/-^)* mice^50^, a motor we showed is required for the anterograde transport of YFP-PrP^C^ vesicles in axons^50^. Depletion of KIF5C resulted in an overall impairment of anterograde vesicle dynamics including a significant reduction in the anterograde flux of PrP^PG14^-EGFP vesicles into the axon, and in decreased vesicle densities (**Fig. 4a-c and Supplementary Fig. 4a**). Moreover, more PrP^PG14^-EGFP vesicles were stationary at the expense of those moving in the anterograde direction (**Fig. 4d**). Vesicles also spent less time moving toward the synapse, moved slower, and paused for longer periods during an anterograde run (**Fig. 4e and Supplementary Fig. 4b,c**). Overall, retrograde PrP^PG14^-EGFP vesicle dynamics were impaired to a lesser extent in *Kif5c^-/-^* axons, and these vesicles spend more time moving toward the soma (**Fig. 4e**). These data indicate that KIF5C drives the anterograde flux of PrP^PG14^-EGFP vesicles from the soma into the axon.

**Figure 4.**
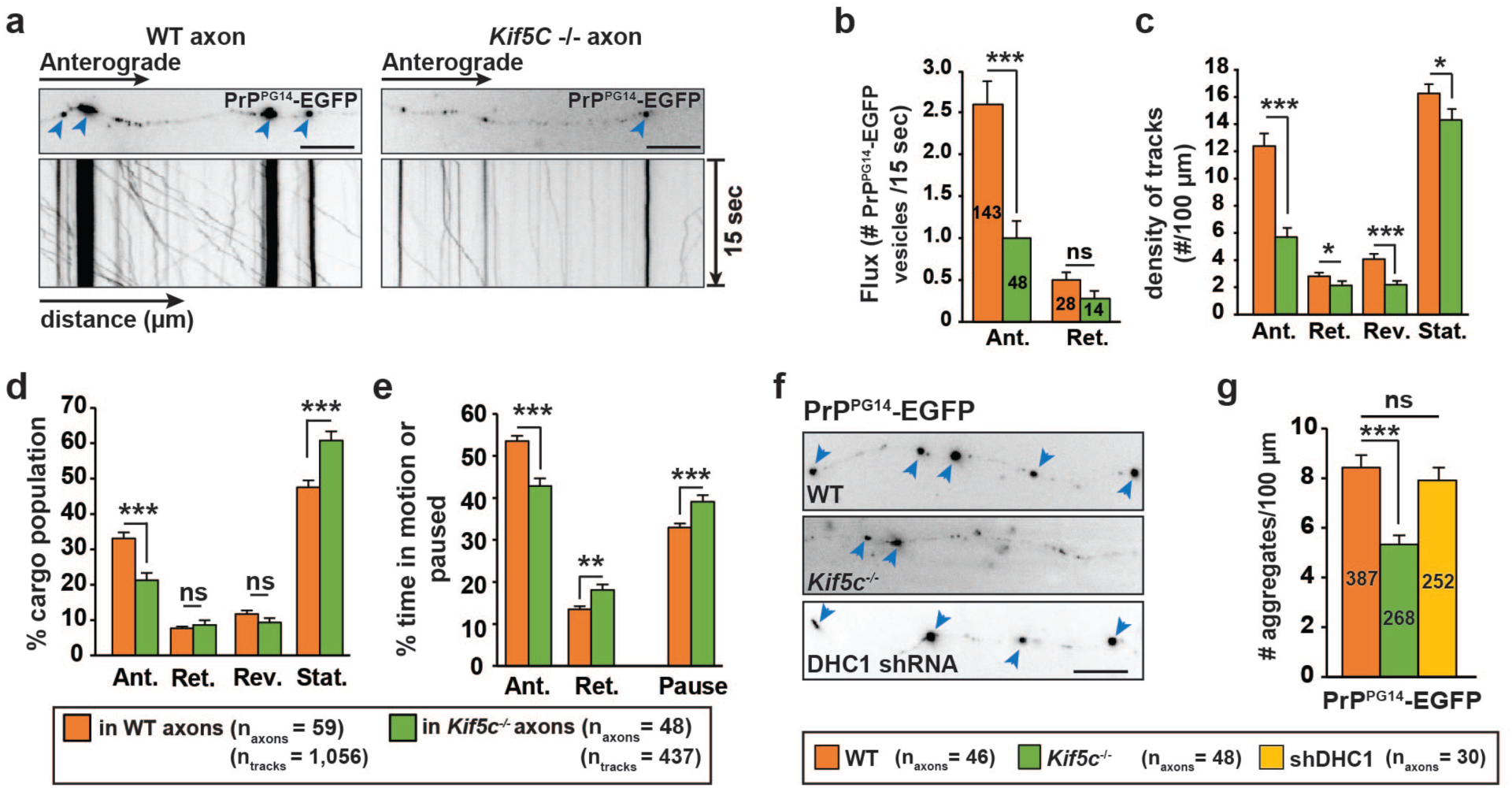
**Kinesin-1 mediates the transport and aggregation of endoggresomes. a**, Representative first frames and kymographs of PrP^PG14^-EGFP vesicle transport or aggregates (arrowheads) in wild-type (WT) (left) and *Kif5c^-/-^* (right) hippocampal axons. Scale bars = 10 μm. **b**, PrP^PG14^-EGFP vesicle flux quantitation. N_vesicles/aggregates_ are inside bars. **c**, Densities of PrP^PG14^-EGFP vesicles in WT or *Kif5c^-/-^* axons. **d**, Population breakdown of PrP^PG14^-EGFP vesicles in WT and *Kif5c^-/-^* axons. **e**, Percent time PrP^PG14^-EGFP vesicles spent in motion or paused in WT and *Kif5c^-/-^* axons. **f**, Representative images of PrP^PG14^-EGFP aggregates (arrowheads) in WT, *Kif5c^-/-^*, and DHC1 shRNA axons. Scale bar = 10 μm. **g**, Quantitation of PrP^PG14^-EGFP aggregate densities from (**f**). N_vesicles or aggregates_ are shown inside bars. Ant. = anterograde, Ret. = retrograde, Rev. = reversal, Stat. = stationary. All values are shown as mean ± SEM. ***p<0.001, **p < 0.01, *p < 0.05, ns = not significant, Student’s t-test. See also **Supplementary Figure 4.**

To next investigate whether active PrP^PG14^-mCh vesicle transport contributed to endoggresome formation, we quantitated PrP^PG14^-mCh aggregate densities in *Kif5c^-/-^* axons. Strikingly, 2 days-post PrP^PG14^-mCh transfection, aggregate densities were significantly decreased in *Kif5c^-^*^/-^ compared to WT hippocampal axons (**Fig. 4f,g**), and these levels were sustained for at least 10 days (**Supplementary Figure 4d,e**). Overexpression of EGFP-KIF5C in *Kif5c^-/-^* neurons transfected with PrP^PG14^-mCh restored higher aggregate densities, showing that the decreased densities were due to the specific removal of KIF5C (**Supplementary Fig. 4f,g**). To further test the role of kinesin-1, we quantitated aggregate densities in axons of cultured hippocampal neurons from kinesin light chain 1 (KLC1) KO (*Klc1^-/-^)* mice, and from conditional kinesin-1B (*Kif5b*) KO mice^72^, both previously implicated in the anterograde transport of YFP-PrP^C^ vesicles^50^. PrP^PG14^-mCh aggregate densities were also significantly decreased in *Klc1*^-/-^ neurons, and in those of conditional *Kif5b* KO neurons in a cre-adenovirus MOI dose-dependent manner (**Supplementary Fig. 4h-l**). Notably, aggregate densities in axons of *Kif5c^-/-^* neurons expressing PrP^D177N(M128)^-mCh were also significantly lower (**Supplementary Fig. 4m,n**), indicating a general requirement of kinesin-1 for generating intra-axonal endoggresomes. To test whether decreased aggregate densities were the result of generalized transport impairments, we reduced the function of the main neuronal retrograde motor cytoplasmic dynein heavy chain 1 (DHC1), using previously validated shRNAs^50^. Despite impairing PrP^PG14^-mCh vesicle transport, and thus identifying DHC1 as a retrograde motor for these vesicles (**Supplementary Fig. 4o**), reducing DHC1 did not alter PrP^PG14^-EGFP aggregate densities (**Fig. 4f,g**). Collectively, these data reveal that in addition to mediating the anterograde flux of PrP^PG14^ vesicles, kinesin-1-mediated anterograde transport is required to form endoggresomes in axons.

### Transient access of misfolded PrP^PG14^ to the cell surface along axons is required for endoggresome formation

To further probe the mechanism of endoggresome formation, we characterized the Golgi-to-endosome sorting itineraries of PrP^PG14^-mCh particles in axons. Specifically, we tested whether upon entering the axon, PrP^PG14^-mCh accessed the plasma membrane *en route* to aggregation. Neurons expressing PrP^PG14^-mCh-BBS were treated with BTX-a647 for 10 minutes at 4°C to inhibit active endocytosis, and then fixed without permeabilization prior to imaging. BTX-a647 signal colocalized with a subset of PrP^PG14^- mCh-BBS-positive puncta along axons (**Fig. 5a**), indicating that the latter accessed the axonal plasma membrane. BTX-a647 labeling was specific, as a647 signal was not observed in neurons expressing PrP^WT^-mCh or PrP^PG14^-mCh without the BBS sequence (**Supplementary Fig. 5a**). To determine if PrP^PG14^ internalized into LEs, we treated PrP^PG14-^mCh-BBS-expressing neurons co-expressing LAMP1-EGFP, with BTX-a647 for 2 hours at 37°C prior to live imaging or fixation and permeabilization. We observed extensive colocalization and cotransport between intra-axonal PrP^PG14-^mCh-BBS, LAMP-1-EGFP puncta/aggregates, and BTX-a647 signal (**Fig. 5b and Supplementary Fig. 5b**), indicating PrP^PG14-^mCh-BBS sorted into LEs post-endocytosis. BTX-a647 signal was observed along axons, demonstrating that PrP^PG14^ undergoes dynamic bouts of exo- and endocytosis throughout the axonal surface. Importantly, addition of the BBS sequence did not alter endoggresome densities in axons (**Supplementary Fig. 5c**). Inhibition of endocytosis with Dynasore, a dynamin GTPase inhibitor^73^ resulted in a significant, albeit partial decrease in internalization of BTX-a647-labeled PrP^PG14^-mCh particles (**Supplementary Fig. 5d**), suggesting that PrP^PG14^-mCh is partly internalized by clathrin-dependent pathways.

**Figure 5.**
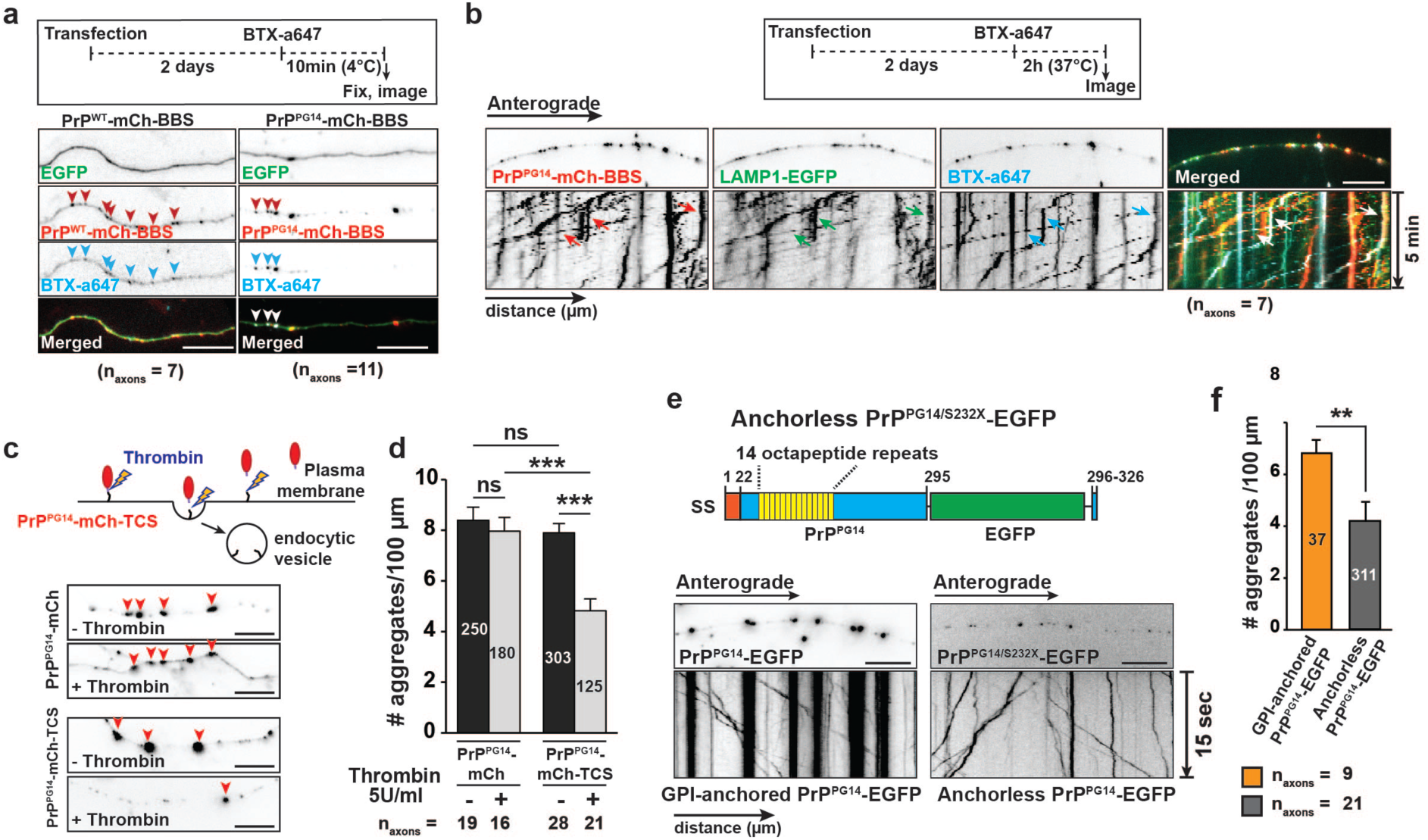
**PrP^PG14^ traffics to the axonal cell surface prior to aggregation. a**, Experimental outline of 10 min BBS internalization assay (top). Representative images of axons expressing EGFP and PrP^WT^-mCh-BBS or PrP^PG14^-mCh-BBS and labeled with BTX-a647 (bottom). Arrowheads point to colocalization. Scale bars = 10 μm. **b**, Experimental outline of 2 hr BBS internalization assay (top). Representative first frames of time-lapse movie, and kymographs of axons expressing PrP^PG14^-mCh-BBS and LAMP1-EGFP, and labeled with BTX-a647. Arrows point to cotransport. Scale bar = 10 μm. **c**, Diagram of thrombin assay (top). Representative images of axons expressing PrP^PG14^-mCh or PrP^PG14^-mCh-TCS, with or without thrombin treatment (bottom). Arrowheads point to aggregates. Scale bars = 10 μm. **d**, Quantitation of aggregate densities from (**c**). N_aggregates_ shown inside bars. **e**, Schematic of anchorless PrP^PG14/S232X^- EGFP construct (top). SS: signal sequence. Representative images and kymographs of hippocampal axons expressing PrP^PG14^-EGFP or PrP^PG14/S232X^-EGFP (bottom). Scale bars = 10 μm. **f**, Quantitation of aggregate densities from (**e**). N_aggregates_ shown inside bars. All values are shown as mean ± SEM. ***p<0.001, **p<0.01, Student’s t-test, Tukey’s correction (**d**). See also **Supplementary Figure 5**.

We next investigated whether PrP^PG14^-mCh cell surface targeting and endocytosis was required to form intra-axonal endoggresomes. We quantitated aggregate densities in axons of neurons expressing PrP^PG14^-mCh tagged or not with a Thrombin Cleavage Sequence (PrP^PG14^-mCh-TCS), and treated or not with 5 units/ml of Thrombin protease for 48 hours to cleave PrP-mCh-TCS at the cell surface prior to endocytosis (**Supplementary Fig. 5e**). Thrombin treatment resulted in significantly reduced densities of PrP^PG14^-mCh-TCS endoggresomes compared to untagged PrP^PG14^-mCh (**Fig. 5c,d**), indicating that access to the plasma membrane is required for their formation. We further tested this cell surface requirement by expressing an anchorless PrP^PG14^ double mutant (PrP^PG14/S232X^-EGFP) that is unable to associate with membranes^47^. While small vesicles carrying this mutant were actively transported in axons, the density of aggregates was largely reduced (**Fig. 5e,f**). Notably, co-expression with GPI-anchored PrP^PG14^-mCh resulted in PrP^PG14/S232X^-EGFP accumulation in large puncta (**Supplementary Fig. 5f**), suggesting that deleting the GPI anchor did not abolish the propensity of this mutant to either misfold and form aggregates, or to accumulate. Altogether, these data indicate that transient cell surface access of PrP^PG14^ occurs along the length of axons, and is required to form PrP^PG14^-mCh endoggresomes in LEs. As endoggresome formation requires transport-dependent sorting (**Fig. 4 and 5**), we term this process **<u>a</u>**xonal **<u>r</u>**apid **<u>e</u>**ndosomal **<u>s</u>**orting and **t**ransport-dependent **<u>a</u>**ggregation (ARESTA), and distinguish it from RESET, which despite involving a similar PrP^PG14^ cell surface-to-lysosome itinerary, has an opposite role in the soma to clear misfolded PrP^PG14^ (**Fig. 1**).

### Arl8b drives ARESTA-mediated formation of axonal PrP^PG14^ endoggresomes via recruitment of kinesin-1 and HOPS

While cell surface ARESTA (henceforth termed “indirect ARESTA”), is required for endoggresome formation (**Fig. 5**), two observations led us to hypothesize that direct Golgi-to-LE or LE homotypic fusion^74–76^ might also contribute to endoggresome biogenesis in axons. First, PrP^PG14^ particles were internalized from the cell surface in single endosomes (**Fig. 5b**), but over a period of 2-5 days, endoggresomes were observed in enlarged LEs (**Fig. 2b and 3e**), suggesting fusion of endosomes post-endocytosis. Second, inhibiting indirect ARESTA reduced but did not abolish the number of PrP^PG14^ endoggresomes (**Fig. 5c,d**), suggesting the presence of parallel aggregate-forming pathways.

As Arl8b orchestrates the trafficking and fusion of LEs^53–56, 77, 78^, we investigated whether it contributed to the formation of aggregates via association with PrP^PG14^ endosomal vesicles to direct their homotypic fusion in axons. We first tested whether Arl8b colocalized with PrP^PG14^ vesicles. Live imaging 2 days post-co-expression of Arl8b^WT^-mCh and PrP^PG14^-mTagBFP2, showed their colocalization in the soma, and in the cotransport of these vesicles and those carrying ss-NPY-EGFP in the axon, suggesting that Arl8b loaded onto post-Golgi PrP^PG14^ vesicles and entered the axon as a complex (**Supplementary Fig. 6a,b**). Endogenous Arl8b as recognized with an anti-Arl8b antibody, also colocalized with PrP^PG14^ vesicles in soma and axons (**Supplementary Fig. S8**). Furthermore, a majority of PrP^PG14^-mCh vesicles co-migrated with Arl8b^WT^-GFP or GTP-bound Arl8b^Q75L^-GFP, and LAMP1-mTagBFP2 particles, but not with soluble dominant negative (DN) GDP-bound Arl8b^T34N^-GFP^53^ (**Fig. 6a,b and Supplementary Fig. 6c**), indicating that GTPase activity is required for Arl8b association with PrP^PG14^-mCh vesicles. To next test the role of Arl8b in aggregation, we quantitated endoggresome densities following Arl8b reduction or overexpression. We observed significantly less aggregates following Arl8b knock-down using shRNAs (**Supplementary Fig. 6d-f**), and densities were lower in neurons expressing untagged GDP-bound Arl8b^T34N^ (**Fig. 6c,d**), indicating that Arl8b is required for PrP^PG14^-mCh aggregation. In contrast, significantly higher endoggresome densities were observed in axons of neurons overexpressing GTP-active untagged Arl8b^WT^ or Arl8b^Q75L^ (**Fig. 6c,d**), demonstrating that membrane-bound Arl8b is sufficient to drive endoggresome formation.

**Figure 6.**
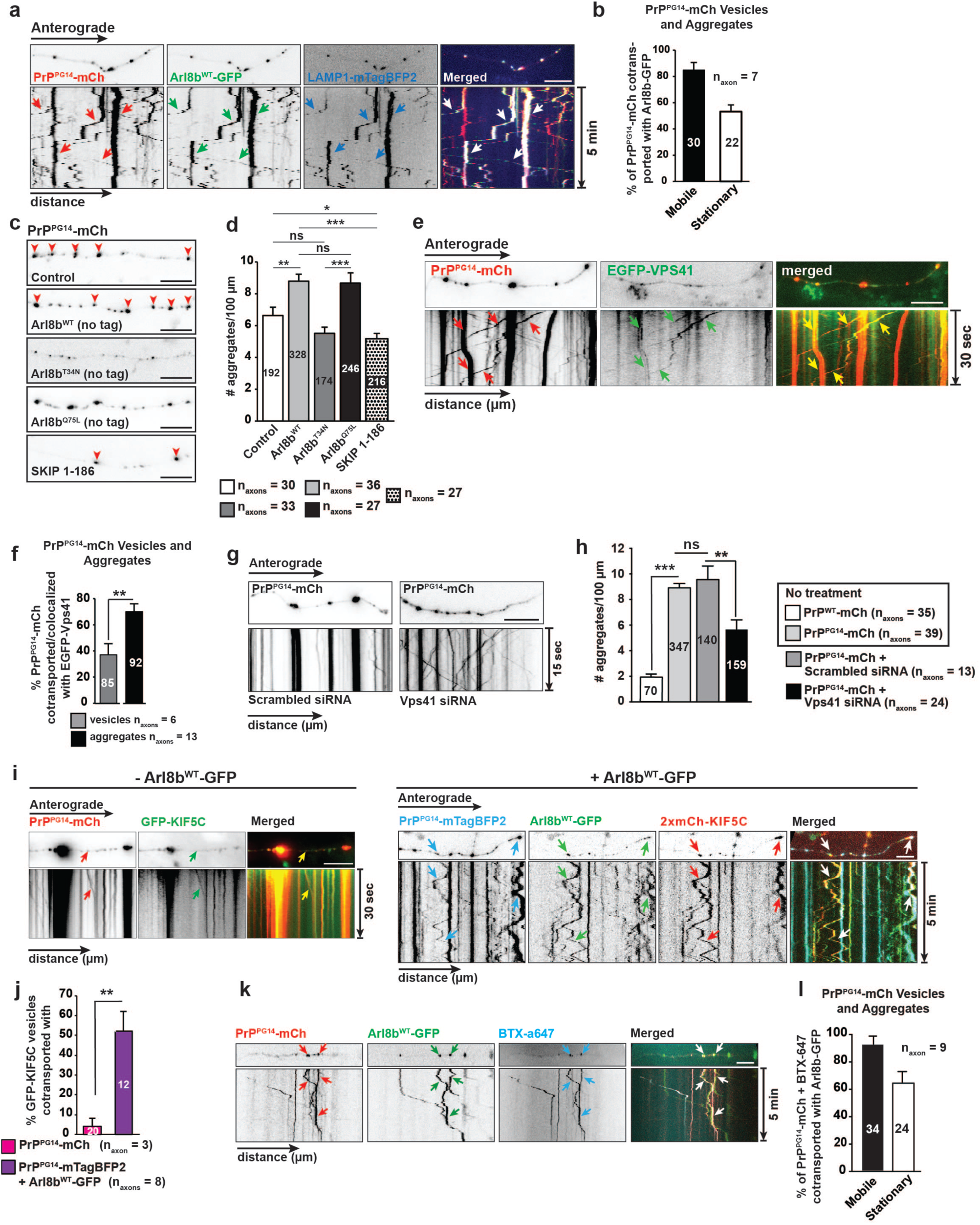
**Arl8b drives PrP^PG14^ aggregation via transport, sorting, and fusion. a**, Representative first frames of time-lapse movie, and kymographs of axons expressing PrP^PG14^-mCh, Arl8b^WT^-EGFP, and LAMP1-mTagBFP2. Arrows point to cotransporting tracks. Scale bar = 10 μm. **b**, Quantitation of cotransport between mobile or stationary PrP^PG14^-mCh vesicles and aggregates and Arl8b^WT^-GFP. N_vesicles/aggregates_ shown inside bars. **c**, Representative images of PrP^PG14^-mCh axons overexpressing empty vector (Control), Arl8b^WT^, dominant-negative (DN) GDP-found Arl8b^T34N^, or GTP-bound Arl8b^Q75L^. Scale bars = 10 μm. **d**, Quantitation of aggregate densities from (**c**) plus aggregate density of axons treated with SKIP 1-186. N_aggregates_ shown inside bars. **e**, Representative first frames of time-lapse movie, and kymographs of axons expressing PrP^PG14^-mCh and EGFP-VPS41. Arrows point to cotransporting tracks. Scale bar = 10 μm. **f**, Quantitation of PrP^PG14^-mCh vesicles and aggregates that cotransport/colocalize with EGFP-VPS41. N_vesicles/aggregates_ are shown inside bars. **g**, Representative first frames of time-lapse movies, and kymographs of axons co-expressing PrP^PG14^-mCh and scrambled or Vps41 siRNAs. Scale bar = 10 μm. **h,** Quantitation of aggregate densities from (**g**). N_aggregates_ are shown inside bars. **i**, Representative first frames of time-lapse movie, and kymographs of axons expressing PrP^PG14^-mCh and GFP-KIF5C (left), and of those expressing PrP^PG14^-mTagBFP2, Arl8b^WT^-GFP, and 2xmCh-KIF5C (right). Arrows point to cotransporting vesicles/aggregates. Scale bars = 10 μm. **j**, Quantitation of cotransport of (**i**). N_vesicles_ are shown inside bars. **k**, Representative first frames of time-lapse movie, and kymographs of axons expressing PrP^PG14^-mCh-BBS, Arl8b^WT^-GFP, and labeled with BTX-a647. Arrows point to col-transporting vesicles. Scale bar = 10 μm. **l**, Quantitation of cotransport of BTX-a647-labeled PrP^PG14^-mCh-BBS vesicles and aggregates with Arl8b^WT^-GFP from (**k**). N_vesicles_ are shown inside bars. All values are shown as mean ± SEM. ***p<0.001, **p < 0.01, *p<0.05, ns = not significant, Student’s t-test (**f**), Šidák correction (**d, h, j**). N_vesicles/aggregates_ shown on graphs. See also Supplementary Figures 6 and 7.

We next examined the mechanistic basis of PrP^PG14^-mCh-Arl8b-mediated aggregate build-up in axons. Arl8b is known to recruit kinesin-1 (KLC1 and KIF5) to LEs via binding of its effector SifA and kinesin-interacting protein (SKIP)^54, 79^, and to directly bind to Vps41, a unit of the HOPS complex required for LE tethering and fusion^80^. We posited that Arl8b drives endoggresome formation by recruiting SKIP, KIF5C, and Vps41 to PrP^PG14^ vesicles, to both propel them anterogradely and to promote their fusion in axons. Live imaging showed extensive cotransport between vesicles carrying PrP^PG14^-mTagBFP2, Arl8b^WT^-mCh, and EGFP-hSKIP (**Supplementary Fig.6g-k**), and between those carrying PrP^PG14^-mCh and EGFP-Vps41 (**Fig. 6e,f**). To investigate the role of Vps41 and SKIP in the formation of endoggresomes, we quantitated aggregate densities in axons of neurons treated either with Vps41 siRNAs (**Supplementary Fig. 6l-o**), or with a SKIP inhibitory peptide that we designed comprising region aa 1-186 (SKIP 1-186), previously shown to be required to bridge interactions between Arl8b and kinesin-1^79^. Aggregate densities were significantly reduced in axons treated with SKIP 1-186 and Vps41 siRNAs (**Fig. 6c,d,g,h**). Furthermore, reducing Vps41 resulted in smaller and less stationary PrP^PG14^-mCh vesicles in the soma and axons, suggesting decreased vesicle fusion and restored motility with less aggregates (**Fig. 6g and Supplementary Fig. 6p,q**).

Next, we tested whether Arl8b recruited kinesin-1 to PrP^PG14^ vesicles to direct their movement into the axon. Overexpression of Arl8b^WT^-mCh resulted in a striking increase in the percentage of PrP^PG14^-mTagBFP2 vesicles and aggregates that cotransported with 2xmCh-KIF5C (**Fig. 6i-j**), and this recruitment depended on Arl8b GTPase activity (**Supplementary Fig. 7a**). Moreover, the flux of PrP^PG14^-mCh vesicles was significantly enhanced in axons overexpressing untagged Arl8b, resulting in increased numbers of moving PrP^PG14^-mCh vesicles into the axon (**Supplementary Fig. 7b-d**). Altogether, these data indicate that GTPase-active Arl8b^WT^ loads onto Golgi-derived PrP^PG14^ vesicles, recruits SKIP, kinesin-1, and Vps41, and acts as a key driver of the movement and direct fusion of PrP^PG14^ vesicles in the axon, thus contributing towards the biogenesis of PrP^PG14^ endoggresomes via a process we termed “direct” ARESTA.

### Direct and indirect ARESTA converge on Arl8b to form axonal endoggresomes

To investigate whether direct and indirect ARESTA act independently or in concert to generate PrP^PG14^ axonal endoggresomes, we overexpressed Arl8b to promote aggregate formation, while blocking indirect ARESTA by inhibiting the uptake of cell surface PrP^PG14^-mCh. Live imaging of neurons expressing PrP^PG14^-mCh-TCS and simultaneously transfected with Arl8b^WT^-GFP at a time point when axons otherwise exhibited few endoggresomes (day 1 post-PrP^PG14^-mCh-TCS transfection), resulted in the same level of increased aggregate densities in axons treated or not with thrombin (**Supplementary Fig. 7e,f**). These observations indicate that Arl8b^WT^-GFP overexpression alone was sufficient to promote endoggresome formation, bypassing the contribution of cell surface PrP^PG14^-mCh, and suggest that direct and indirect ARESTA act in parallel.

We next investigated whether Arl8b associated with PrP^PG14^ vesicles endocytosed from the cell surface following indirect ARESTA. Live imaging of axons expressing PrP^PG14^-mCh-BBS, Arl8b^WT^-GFP, and treated in the media with BTX-a647 resulted in significant cotransport between these three vesicle populations (**Fig. 6k,l**), indicating Arl8b can intersect endocytosed PrP^PG14^-mCh-BBS vesicles. Collectively, these data indicate that direct and indirect ARESTA act in parallel and converge on Arl8b to drive the formation of axonal PrP^PG14^ aggregates. These observations position Arl8b as a key regulator of both the direct fusion of Golgi-to-LE PrP^PG14^ vesicles, as well as of the endosomal sorting of PrP^PG14^ internalized from the cell surface into endosomes that drives the biogenesis of mutant prion endoggresomes in axons.

### Arl8b is a key determinant of PrP^PG14^ axonal entry

As Arl8b is an endosomal GTPase, we tested whether it associated uniquely with post-Golgi endosomal PrP^PG14^ vesicles destined for axonal entry, thus distinguishing these from PrP^PG14^ vesicles secreted in the soma for RESET degradation (**Fig. 1**). To test this, we followed the ER-to-Golgi-to-LE secretory itinerary of PrP^PG14^-mTagBFP2 (or -mCh) and Arl8b in the soma in neurons treated or not with BFA. To prevent overexpression of Arl8b, we analyzed the localization of endogenous Arl8b in fixed cells using a previously validated antibody^4^. Colocalization analysis showed that as observed earlier (**Fig. 1**), PrP^PG14^-mTagBFP2 (or -mCh) translocated from the ER to the Golgi, and to LAMP1-mTag FBP2-positive endo-lysosomes immediately post-BFA wash (**Supplementary Fig. 8a**). However, Arl8b did not extensively colocalize in the soma with PrP^PG14^-mTagBFP2 throughout this itinerary, although some colocalization was observed suggesting Arl8b started association with PrP^PG14^ vesicles 2 hrs post-transfection (**Supplementary Fig. 8a, i, ii**). Instead, colocalization of Arl8b with PrP^PG14^-mCh and LAMP1-mTagBFP2 vesicles occurred primarily in the proximal axon, as observed earlier in more distal axonal regions (**Fig. 6**). As Arl8b associates in the axon with PrP^PG14^ vesicles that directly endocytose from the cell surface (**Fig. 6k,j**), we investigated whether Arl8b also associated with PrP^PG14^ vesicles that internalized from the somatic cell surface following RESET. Immunofluorescence of neurons co-transfected with PrP^PG14^-mCh-BBS and LAMP1-mTagBFP2 and stained with an antibody against Arl8b revealed the colocalization of PrP^PG14^-mCh-BBS and LAMP1-mTagBFP2 vesicles that were also positive for BTX-a647 signal, but that were negative for Arl8b (**Supplementary Fig. 8b**) indicating that unlike in the axon, Arl8b does not recognize PrP^PG14^ vesicles that are internalized from the somatic cell surface. Collectively, these data show that Arl8 does not associate with somatic PrP^PG14^ vesicles undergoing RESET, but instead uniquely loads onto post-Golgi PrP^PG14^ vesicles that enter the axon, suggesting that Arl8b is a key regulator of PrP^PG14^ axonal entry.

### Impaired retrograde transport and lysosomal degradation sustain endoggresomes in axons

We next investigated how PrP^PG14^ endoggresomes persisted in axons. Previous studies revealed that while limited, the axon has some local degradation capacity^4^, although the bulk of degradation of axonal cargoes occurs primarily via their retrograde long-distance movement back to the soma for fusion with degradative lysosomes^5^. Thus, we tested whether persistence of PrP^PG14^ endoggresomes in axons was due to impairments in local axonal lysosomal degradation capacity, and/or failure of PrP^PG14^ vesicles to undergo retrograde transport. Several experiments suggested deficits in endo-lysosomal maturation and lack of local active axonal PrP^PG14^ degradation. First, axonal PrP^PG14^-mCh-positive aggregate densities increased over time compared to PrP^WT^-mCh-positive puncta (**Fig. 2b,c**), suggesting impaired *in situ* degradation. Additionally, treatment of PrP^PG14^-mCh-expressing neurons with Bafilomycin A1 (BafA1), an inhibitor of a lysosomal V-ATPase, a proton pump that acidifies vesicles^81^, did not increase the density of aggregates compared to DMSO-treated controls **(Supplementary Fig. 9a**). As a positive control, PrP^WT^-mCh neurons treated with BafA1 showed a significant increase in aggregate densities compared to DMSO-treated ones, indicating inhibition of acidification (**Supplementary Fig. 9a**). These data suggest that PrP^PG14^ aggregates reside in endo-lysosomal compartments that are not actively degradative. Consistent with this observation, and in contrast to the soma (**Fig. 1d**), LysoTracker signal was not readily observed in PrP^PG14^-mCh axons (**Fig. 7a,b and Supplementary Fig. 9b**), showing compromised lysosomal degradative potential. Third, neurons expressing PrP^PG14^-mCh-BBS showed particles that were actively transported inside axons 24 hours following BTX-a647 treatment and wash-off, indicating that internalized PrP^PG14^-mCh-BBS molecules were not degraded in lysosomes (**Fig. 7c**).

**Figure 7.**
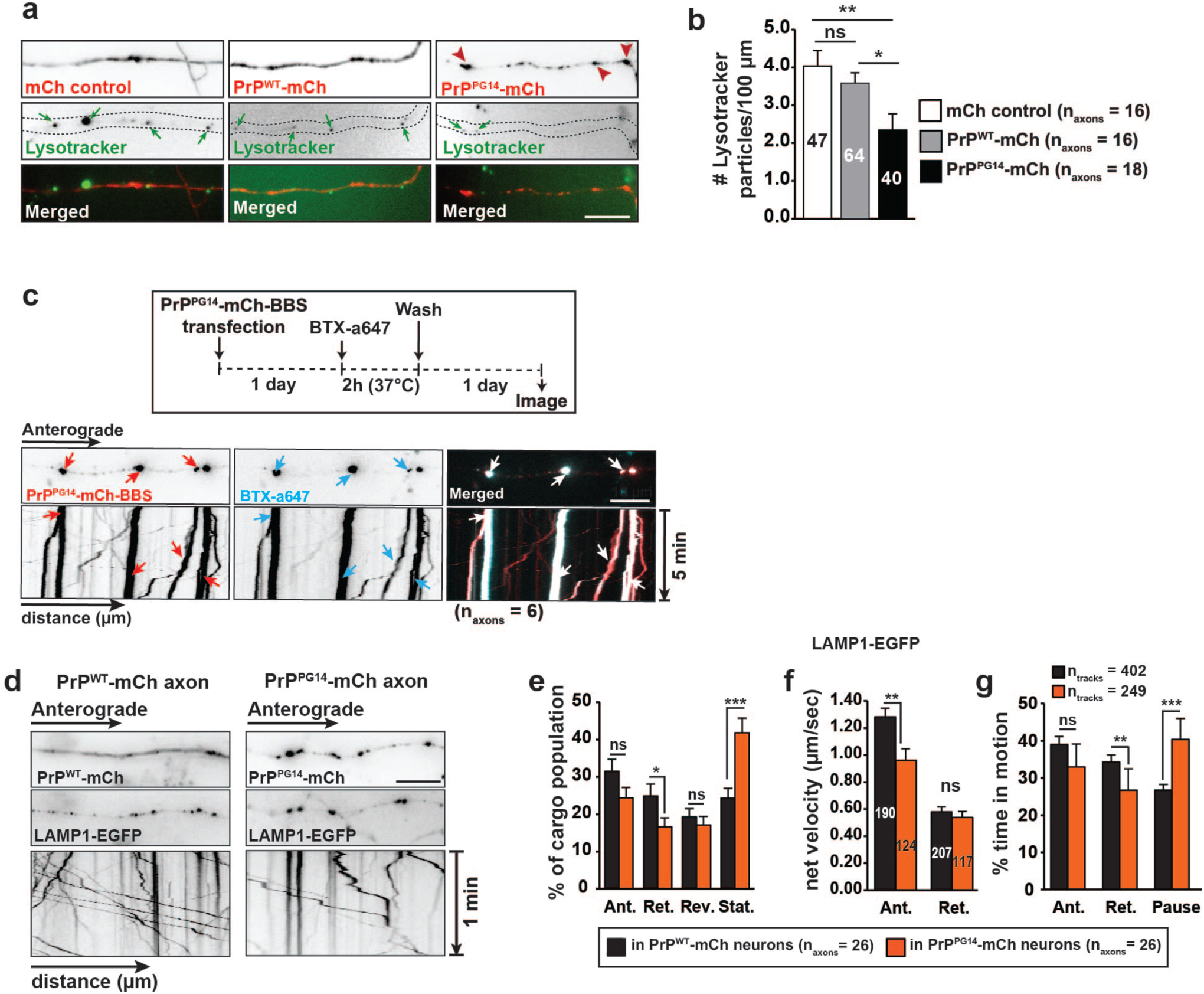
**Impaired lysosomal maturation and retrograde transport of PrP^PG14^ endoggresomes. a**, Representative images of Lysotracker-positive vesicles (arrows) in mCh control, PrP^WT^-mCh, or PrP^PG14^-mCh-expressing axons. Arrowheads point to aggregates. Scale bar = 10 μm. **b**, Quantitation of Lysotracker vesicle densities from (**a**). N_vesicles_ are shown inside bars. **c**, Experimental outline of BTX assay (top). Representative first frames of time-lapse movie, and kymographs of BTX-a647-labeled PrP^PG14^-mCh-BBS axons. Arrows point to cotransporting tracks. Scale bar = 10 μm. **d**, Representative images of PrP^WT^-mCh or PrP^PG14^-mCh axons (top), and LAMP1-EGFP first frames of time-lapse movie (middle), and kymographs (bottom). Scale bar = 10 μm. **e**, Population breakdown of LAMP1-EGFP vesicles in PrP^WT^-mCh or PrP^PG14^-mCh axons. **f**, Average net velocities of LAMP1-EGFP vesicles in PrP^WT^-mCh or PrP^PG14^-mCh axons. N_tracks_ are shown inside bars. **g**, Percent time LAMP1-EGFP vesicles spent in motion or paused in PrP^WT^-mCh or PrP^PG14^-mCh axons. All values are shown as mean ± SEM. ***p<0.001, **p < 0.01, *p<0.05, ns = not significant, Student’s t-test (**e-g**), Tukey’s correction (**b**). See also **Supplementary Figure 9.**

We next tested whether the retrograde transport of PrP^PG14^ vesicles was impaired. In addition to the largely stationary or slow moving PrP^PG14^-mCh endoggresomes (**Fig. 3a,b**), live imaging and quantitative analysis showed PrP^PG14^-EGFP vesicles also moved significantly slower and with less processivity in the anterograde and retrograde directions compared to PrP^WT^-EGFP vesicles (**Supplementary Fig. 9c**). To further test whether retrograde transport was impaired, we imaged the transport of LAMP1-EGFP vesicles in PrP^WT^-mCh versus PrP^PG14^-mCh expressing neurons. The proportion of stationary LAMP1-EGFP particles in PrP^PG14^ axons was significantly increased, and less vesicles traveled retrogradely, moved slower in the anterograde direction, and spent more time paused and less time moving toward the soma (**Fig. 7d-g**). These results show defective retrograde and anterograde movement of LAMP1-EGFP vesicles, suggesting impaired distribution of axonal LEs in PrP^PG14^ axons. Collectively, our data indicate that PrP^PG14^ vesicles and endoggresomes, as well as LEs are not properly transported toward the soma, nor are they actively degraded in axonal lysosomes, contributing to sustained presence of PrP^PG14^ endoggresomes in axons.

### Functional consequences of kinesin-1-dependent formation of PrP^PG14^ endoggresomes in axons

To probe whether axonal PrP^PG14^ endoggresomes lead to neuronal dysfunction, we tested whether the intrinsic capacity of cultured hippocampal neurons expressing PrP^PG14^-mCh to intake calcium was compromised upon KCl-induced depolarization, as shown previously for cerebellar granule culture neurons^82^. Cultured primary hippocampal neurons were transduced with PrP^WT^-mCh or PrP^PG14^-mCh Adeno-associated virus (AAV-DJ), to obtain high transduction efficiencies of 96% ± 2% and 97% ± 2% (mean ± SEM), respectively. Comparable endoggresome densities were observed in transduced versus transfected axons (**Supplementary Fig. 10d**). Single-cell calcium imaging of AAV-DJ-PrP^PG14^-mCh-transduced neurons pre-loaded with the calcium sensitive dye Fluo-4 AM resulted in a defective calcium influx in response to KCl-induced depolarization compared to neurons transduced with AAV-DJ-PrP^WT^-mCh (**Fig. 8a-c**). We investigated whether reducing endoggresome formation via depletion of ARESTA component kinesin-1 could inhibit calcium intake dynamics. Strikingly, *Kif5c^-/-^* neurons with significantly less endoggresomes (**Fig. 4f,g**), but treated with AAV-DJ-PrP^PG14^-mCh displayed normal calcium influx dynamics comparable to those of neurons expressing AAV-DJ-PrP^WT^-mCh (**Fig. 8a-c**). Importantly, calcium intake was not altered in control *Kif5c^-/-^* neurons transduced with AAV-DJ-PrP^WT^-mCh (**Fig. 8d-f**).

**Figure 8.**
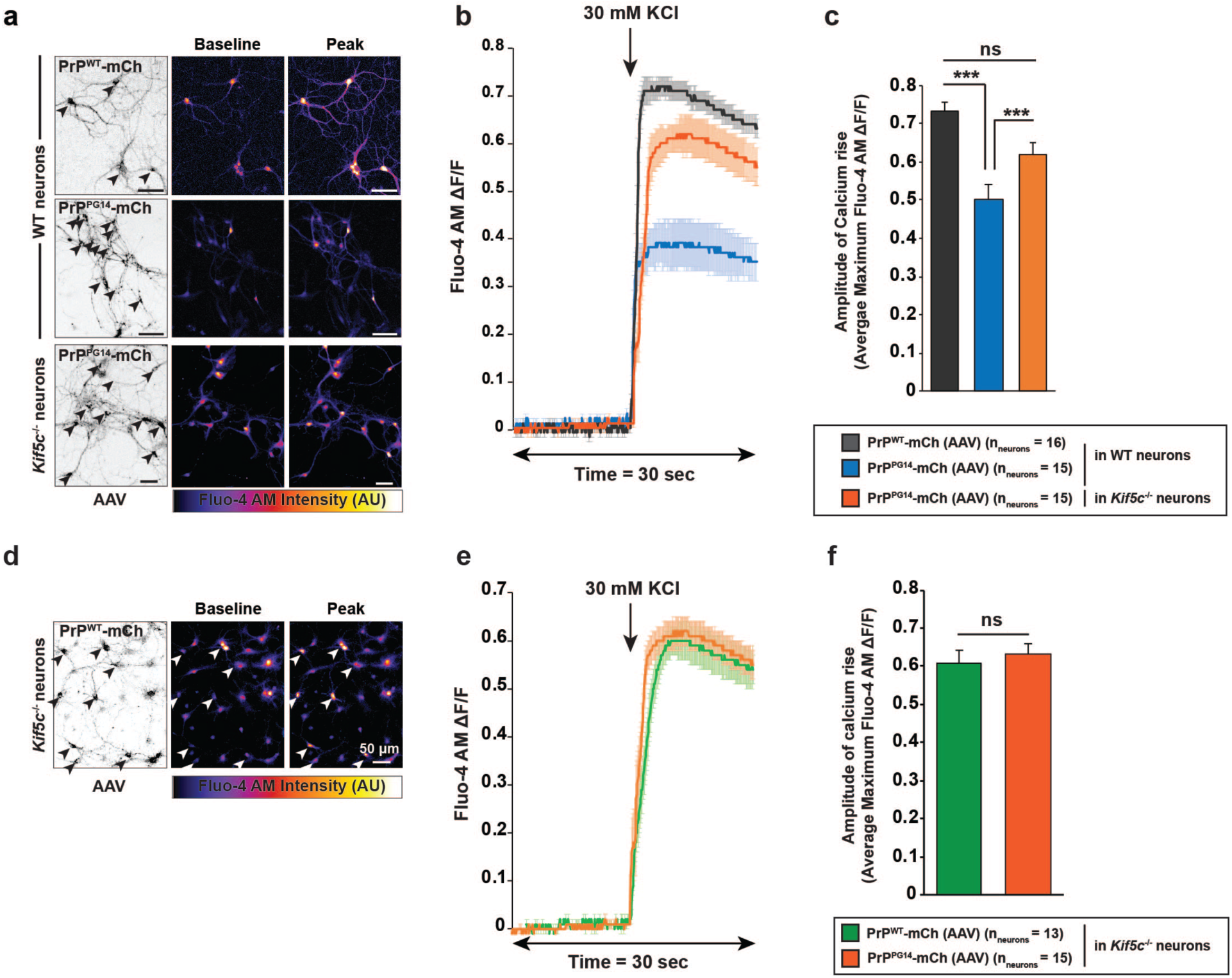
**Kinesin-1-dependent PrP^PG14^ endoggresome formation impairs calcium influx dynamics. a**, Representative images of PrP^WT^-mCh or PrP^PG14^-mCh AAV-transduced WT or *Kif5c*^-/-^ neurons (arrowheads; left). Baseline and peak Fluo-4 AM intensities pre- and post-KCl treatment (middle, right). Scale bars 50 μm. **b**, KCl-induced depolarization response curve from normalized Fluo-4 AM intensities from (**a**). **c**, Average maximum amplitude of calcium rise from (**b**). **d**, Representative images of PrP^WT^-mCh AAV-transduced *Kif5c*^-/-^ neurons (arrowheads). Baseline and peak Fluo-4 AM intensities pre- and post-KCl treatment (middle, right). **e**, KCl-induced depolarization response curve from normalized Fluo-4 AM intensities from (**d** and **a**). **f,** Average maximum amplitude of calcium rise from (**e**). All values are shown as mean ± SEM. ***p<0.001, ns = not significant, Student’s t-test, Tukey’s correction (**c,f**). See also **Supplementary Figure 10.**

To further characterize the effect of PrP^PG14^ endoggresomes, we quantitated cell death in WT versus *Kif5c^-/-^* primary hippocampal neurons. At 10-day post-PrP^PG14^-mCh transfection, when endoggresomes are prominent in WT axons (**Supplementary Fig. 4d,e**), expression of PrP^PG14^-mCh resulted in a ∼4.5 -fold increase in cell death compared to non-transfected neurons or those expressing PrP^WT^-mCh (**Supplementary Fig. 10b,c**). Notably, neuronal death at 10 days post-transfection was not significantly different in *Kif5c^-/-^* neurons expressing PrP^PG14^-mCh compared to those expressing PrP^WT^-mCh or not transfected, suggesting that reduced endoggresome formation in KIF5C-depleted axons prevented cell death. Altogether, our data indicate that the presence of endoggresomes imparts toxicity and affects neuronal viability, and that their inhibition by depletion of kinesin-1 abrogates these defects to prevent neuronal dysfunction and cell death.

## DISCUSSION

By characterizing the trafficking itineraries of a misfolded GPI-anchored PrP mutant in mammalian neurons, we uncovered novel endo-lysosomal pathways that either sort mutant PrP for immediate degradation within lysosomes in the soma in a process that resembles non-neuronal RESET^47, 48^, or that direct mutant PrP into the axon for aggregation via ARESTA inside late endosomal structures which we call endoggresomes. (**Supplementary Fig. 11**). We uncovered the mechanistic basis of selective endoggresome formation in axons: the small GTPase Arl8b associates with post-Golgi vesicles harboring misfolded mutant PrP, earmarking them for axonal entry. Arl8b recruits kinesin-1 and Vps41(HOPS) and this endosomal vesicle-protein complex moves in the anterograde direction along the axon to undergo homotypic fusion and form aggregates. Endoggresomes persist in axons in response to impaired retrograde transport and decreased axonal degradative capacity, resulting in calcium intake defects and accelerated neuronal death. Remarkably, these defects can be circumvented by inhibiting endoggresome formation through the depletion of the ARESTA component kinesin-1 (**Supplementary Fig. 11**).

Our findings have a number of important implications for the understanding of quality control mechanisms that neurons have adopted in response to aggregation-prone proteins residing within the secretory pathway. The discovery of a partitioning itinerary for mutant PrP for somatic degradation versus axonal aggregation, was unexpected. In non-neuronal RESET, essentially all misfolded PrP particles were shown to be degraded in lysosomes via a plasma membrane route, indicating this is a *bone fide* and effective clearance mechanism^48^. In neurons this does not appear to be the case. While the precise ratio of PrP^PG14^ undergoing RESET versus ARESTA is unknown, our data argue that mutant PrP can escape immediate degradation and sort into axons. We posit that in neurons, segregation toward degradation via RESET could be triggered by ER stress activation, as proposed for non-neuronal cells^48^. Under this scenario, non-neuronal misfolded PrP was shown to associate with ER cargo receptors and chaperones, which were proposed to tag the complex for travel to the plasma membrane prior to internalization in lysosomes. Our work shows that expression of PrP^PG14^ robustly activates neuronal ER stress, followed by ER-to-Golgi egress of PrP^PG14^ within minutes, and subsequent targeting to degradative lysosomes via the cell surface within an hour (**Fig. 1**). These dynamics are consistent with non-neuronal RESET, suggesting ER stress could also guide mutant PrP toward secretion and clearance.

Our data provides evidence that sorting of mutant PrP into the axon occurs at the level of the trans-Golgi network (TGN), by engagement of an Arl8b/kinesin/HOPS complex that we propose represents a signature for axonal entry. Arl8b was absent from RESET-bound PrP^PG14^ vesicles, but associated with over 80% of PrP^PG14^ vesicles entering the axon, suggesting it represents an axonal tag (**Fig. 6**). Evidence for a Golgi-to-axon itinerary was also provided by the observation that PrP^PG14^ vesicles co-transport with Arl8b and Golgi-derived markers into the axon, indicating TGN-to-endosome sorting. Both Arl8b and Vps41 have been implicated in directing TGN-to-LE itineraries^55^, so these roles appear to be conserved during their interactions with mutant PrP vesicles. How Arl8b recognizes mutant PrP vesicles at the TGN requires further investigation, but the mechanism could relate to the regulation of endosomal sorting. A plausible participant is BORC (BLOC-One-Related Complex), an Arl8b adaptor that has been shown to couple LEs to Arl8b to regulate their distribution towards the periphery of cells uniquely in axons and not in dendrites^68, 83^. Cellular stress is known to trigger a switch from a BORC-Arl8b to a BORC-LAMTOR (late endosomal/lysosomal adaptor and MAPK and mechanistic target of rapamycin [mTOR] activator) association, resulting in the perinuclear retention of axonal LEs^84^. As such, an attractive possibility is that ER stress activation resulting from mutant PrP expression could delineate the choice between the formation of either of these complexes on PrP^PG14^ vesicles at the TGN, to partition them to somatic versus axonal itineraries.

Once in the axon, mutant PrP undergoes further sorting with Arl8b at the helm. The Arl8b/SKIP/kinesin-1/Vps41 complex provides the axon with a steady flux of mutant PrP vesicles that either fuse with the axonal plasma membrane allowing mutant PrP transient access before internalization in LEs (indirect ARESTA), or directly fuse with other LEs (direct ARESTA) (**Fig. 5, 6**). These parallel pathways converge on Arl8b to drive endoggresome formation. We posit that the short but dynamic residence time of mutant PrP at the axonal cell surface tips the balance towards fusion by increasing the number of PrP^PG14^ vesicles inside axons available to Arl8b for fusion. In contrast, the steady presence of PrP^WT^ at the plasma membrane (**Fig. 2a**) allows for recycling or clearance cycles that balance axonal influx and degradation resulting in an aggregate-free axon.

The identification of ARESTA highlights the discovery of a new type of intracellular aggregate compartment which we call the endoggresome. We define endoggresomes as structures composed of misfolded proteins that are formed inside endomembranes via the action of kinesin-dependent anterograde transport and requiring endosomal fusion, and that are not easily degraded. Previous observations of accumulations of misfolded proteins, including of PrP^Sc^, into cytosolic aggresomes, IPODS, and/or JUNQ structures^85, 86^ rendered our discovery somewhat unexpected. PrP^Sc^ aggresomes were previously observed to form following proteasome inhibition and after retrotranslocation to the cytosol, where they are targeted for clearance by ERAD^37^. However, extensive characterization of the trafficking and turnover of many misfolded GPI-anchored PrP mutants showed that they are normally poor ERAD substrates, are not dependent on proteasomal degradation, and exit the ER for degradation in lysosomes^87^. Thus, it is possible that endoggresome formation in axons is favored by misfolded PrP at steady state, but this may shift towards aggresome formation in the soma when and if RESET is overwhelmed or compromised during ER stress.

Several features distinguish endoggresomes from previously described aggresomes, IPODs or JUNQs^85^. First, the latter three are membrane-less structures while endoggresomes are formed inside endomembranes. Second, cytosolic aggregates depend on dynein-mediated retrograde movement to form at juxtanuclear or perivacuolar sites, but endoggresomes depends on kinesin-based anterograde transport to form peripherally, as removal of kinesin-1 subunits dramatically inhibited their formation (**Fig. 3**). Third, the cores of cytosolic aggresomes, IPODs, and JUNQs are targeted for clearance by autophagosomes and/or proteasomal degradation, but our data show that endoggresomes are not easily targeted for degradation, as shown by the lack of lysosomal clearance that leads to their prolonged presence in axons. Finally, ample evidence suggests that the formation of cytosolic aggregates is a cytoprotective response mechanism to sequester proteins with chaperones to restore their native conformation, or if this fails, to rapidly destroy misfolded protein aggregates^8^. In contrast, our data show that endoggresomes impart toxicity by impairing the ability of neurons to regulate calcium intake dynamics, accelerating neuronal death.

The generality of endoggresomes and ARESTA within the prionopathies and to other neurodegenerative pathologies requires further investigation. However, our analysis indicates that at least in the case of PrP^D178N/M129^, the formation of intra-axonal aggregates also depends on KIF5C (**Supplementary Fig. 3**), suggesting that ARESTA might be a common aggregation mechanism of familial PrP mutations. Given the reliance of tau, APP/A*β*, *α*-synuclein, and huntingtin on endosomal pathways for their processing and spread, it is possible that ARESTA may also be implicated in the formation of misfolded aggregates of these proteins^88–90^. As genetic modulation of ARESTA components resulted in changes in endoggresome densities with measurable functional consequences to the heath and viability of neurons, the identification of the Arl8b/kinesin-1/HOPS ARESTA pathway and of endoggresomes provides an appealing anti-aggregation target that can modulate neuronal dysfunction and delineate the differential vulnerability of axons to aggregate-induced pathologies.

## METHODS

### Mouse Lines and Maintenance

Our mouse protocols were reviewed and approved by Institutional Animal Care and Use Committee (IACUC) at The Scripps Research Institute (Scripps Research) and all colonies were maintained following the guidelines recommended by the Department of Animal Resources (DAR) at Scripps Research. C57BL/6 wild-type mice were obtained from Charles River Laboratories. Kinesin-1C null (*Kif5c*^-/-^) and KLC1 null (*Klc1*^-/-^) mice were generated in Larry Goldstein’s lab^50^. *Kif5c*^-/-^ mice were maintained as homozygotes and were backcrossed in the C57BL/6 background for at least 6 generations. *Klc1*^-/-^ mice were bred as heterozygotes to generate homozygous pups. The conditional homozygous kinesin-1B floxed mouse (*Kif5b^pflox/pflox^*) was generated by Nancy Jenkins and JianDong Huang^72^.

### Primary Neuronal Culture

Primary hippocampal neuron cultures were prepared as described previously^50^. Briefly, hippocampi were dissected from 0 to 2-day-old mouse neonates in cold Hank’s buffer solution (HBSS) (Gibco) supplemented with 0.08% D-glucose (Sigma), 0.17% HEPES, and 1% Pen-Strep (Gibco), filter sterilized and adjusted to pH 7.3. Dissected hippocampi were washed twice with cold HBSS (Gibco) and individually incubated at 37°C for 15-20 minutes in a sterile solution of 45 U papain (Worthington), 0.01% DNase (Sigma), 1 mg DL-cysteine (Sigma), 1 mg BSA (Sigma) and 25 mg D-Glucose (Sigma) in PBS (Gibco). Hippocampi were washed twice with Dulbecco’s Modified Eagle Medium (DMEM) (Gibco) supplemented with 10% fetal bovine serum (FBS) (Gibco) (pre-heated to 37°C) and disrupted by ten to twelve cycles of aspiration through a micropipette tip. Dissociated neurons were then resuspended in warm DMEM (Gibco) + 10% FBS (Gibco) and plated in 24-well plates containing 12-mm glass coverslips pretreated with 50 µg/ml poly-L-lysine (Sigma) in Borate buffer (1.24 g boric acid (Fisher), 1.90 g borax (Sigma), in 500 mL cell-culture grade water, adjusted to pH 8.5, filtered sterilized). After 1 hour, media was replaced with Neurobasal-A medium (NBA) (Gibco), supplemented with 2% B-27 (Gibco) and 0.25% GlutaMAX (Gibco). Primary neurons were maintained in an incubator at 37°C and in a 5.5% CO_2_ atmosphere.

For quantitative live imaging of axonal transport, hippocampal neurons were plated in microfluidic chambers, or in regular 24-well glass coverslips as described previously ^50^. For calcium imaging, hippocampal neurons were plated directly onto #1.5 glass-bottomed dishes (MatTek) precoated with 50 µg/ml poly-L-lysine (Sigma) in Borate buffer.

### Cell Culture

Neuro2a (N2a) cells (ATCC) were grown in DMEM (Gibco), supplemented with 10% FBS (Gibco) and 1% Pen-Strep (Gibco) in an incubator at 37°C and in a 5.5% CO_2_ atmosphere. Cells were passaged twice a week to low confluency using 0.05% Trypsin-EDTA (Gibco).

### Design of DNA constructs

The MoPrP.Xho PrP^WT^-EGFP and MoPrP.Xho PrP^PG14^-EGFP constructs were a gift from David Harris (Boston University)^36, 60^. All other constructs were generated using the In-Fusion^®^ HD Cloning Plus Kit (Clontech) for the final cloning of DNA fragments into the recipient vector. All PCR amplifications were performed using the Phusion^®^ High-Fidelity DNA Polymerase (NEB).

For the construction of MoPrP.Xho PrP^WT^-mCh and MoPrP.Xho PrP^PG14^-mCh constructs, EGFP was replaced with mCh using assembly PCR. Briefly, three individual PCRs were performed to amplify the (1) N-terminus of PrP^WT^ or PrP^PG14^, (2) C-terminus of PrP, (3) mCh including 20 and 22 bp overlaps to the 3’ of (1) and 5’ of (2), respectively.

(1) The N-terminus of PrP was amplified using primers

P1 N-Ter For: 5’-CTAGTGGTACCTCGAGATGGCGAACCTTGGCTAC-3’ P2 N-Ter Rev: 5’-CATGGTGGCGACCGGTGGATCC-3’

(2) The C-terminus of PrP was amplified using primers

P3 C-Ter For: 5’-TCCGGACTCAGATCTCGAGCTCAAG-3’

P4 C-Ter Rev: 5’-AGCAGGAAGGCTCGAGTCATCCCACGATCAGGAA-3’

(3) mCh was amplified using primers

P5 mCh For: 5’-ATCCACCGGTCGCCACCATGGTGAGCAAGGGCGAGGA-3’

P6 mCh Rev: 5’-GCTCGAGATCTGAGTCCGGACTTGTACAGCTCGTCCATGCCG-3’.

PCR products (1-3) were gel purified and subsequently stitched together (1 ng each), using a PCR reaction performed without primers. After 20 cycles, PCR was interrupted, primers P1 For and P4 Rev (0.5 µM each) were added to specifically amplify the complete product. Reaction was supplemented with 100 µM dNTP and 0.5 mM MgCl_2_ and ran for 30 more cycles. Generation of MoPrP.Xho PrP^WT^-mTagBFP2 and MoPrP.Xho PrP^PG14^-mTagBFP2 constructs was done using the same assembly PCR strategy.

P5 mTagBFP2 For: 5’-ATCCACCGGTCGCCACCATGGTGTCTAAGGGCGAAGA-3’

P6 mTagBFP2 Rev: 5’-GCTCGAGATCTGAGTCCGGAATTAAGCTTGTGCCCCAGTT-3’

Generation of the untagged MoPrP.Xho PrP^WT^ and MoPrP.Xho PrP^PG14^ was done using the same assembly PCR strategy.

(1) The N-terminus of PrP was amplified using primers

P1 N-Ter For: 5’-CTAGTGGTACCTCGAGATGGCGAACCTTGGCTAC-3’ P2 N-Ter Rev: 5’-CATGGTGGCGACCGGTGGATCC-3’

(2) The C-terminus of PrP was amplified using primers

P3 C-Ter For: 5’-CCGGTCGCCACCATGTCCGGACTCAGATCTCGAGCTCAAG -3’ P4 C-Ter Rev: 5’-AGCAGGAAGGCTCGAGTCATCCCACGATCAGGAA-3’

Generation of MoPrP.Xho PrP^WT^-PAmCh and MoPrP.Xho PrP^PG14^-PAmCh was done using the same assembly PCR strategy from PAmCh1-C1 (a gift from Vladislav Verkhusha, Albert Einstein College of Medicine, New York)^91^, which contains the following mutations: E26V/A58T/K69N/L84F/N99K/S148L/I165V/Q167P/L169V/I203R. P5 PAmCh For: 5’-ATCCACCGGTCGCCACCATGGTGAGCAAGGGCGAGGA-3’ P6 PAmCh Rev: 5’-GCTCGAGATCTGAGTCCGGACTTGTACAGCTCGTCCATGCCG-3’.

To generate Bungarotoxin binding sequence (BBS)-tagged constructs, MoPrP.Xho PrP^WT^-mCh-BBS and MoPrP.Xho PrP^PG14^-mCh-BBS, a 13 amino-acid BBS tag (WRYYESSLEPYPD) was inserted at the BspEI site in the linker between mCh and C-terminal PrP sequence downstream of mCh. The following oligos were used to amplify the BBS sequence:

BBS For: 5’-CCGGATGGAGATACTACGAGAGCTCCCTGGAGCCCTACCCTGACT-3’

BBS Rev: 5’-TACCTCTATGATGCTCTCGAGGGACCTCGGGATGGGACTGAGGCC-3’

The PCR products were annealed and cloned in PrP^WT^-mCh in pcDNA3.1 plasmid and PrP^PG14^-mCh in pcDNA3.1 plasmid. PrP^WT^-mCh-BBS and PrP^PG14^-mCh-BBS sequences were amplified by PCR and subcloned into the MoPrP.Xho vector. A similar strategy was used to generate the Thrombin cleavage sequence (TCS) tagged constructs MoPrP.Xho PrP^WT^-mCh-TCS and MoPrP.Xho PrP^PG14^-mCh-TCS. The following oligos were used to amplify the 6 amino-acid TCS tag (LVPRGS), and to insert it at the BspEI site downstream of mCh:

TCS For: 5’-GCTGTACAAGTCCGGACTGGTGCCGCGCGGCAGCTCCGGACTCAGATCTC-3’

TCS Rev: 5’-GAGATCTGAGTCCGGAGCTGCCGCGCGGCACCAGTCCGGACTTGTACAGC-3’

To generate the LAMP1-EGFP construct, RFP was replaced with EGFP in the plasmid LAMP1-RFP (a gift from Erika Holzbaur, University of Pennsylvania)^92^. To generate ss(NPY)-EGFP, the first 84 nucleotides of Neuropeptide Y were amplified from mouse brain cDNA and cloned between BglII and BamHI sites in pEGFP-N3 vector. To design EGFP-Rab5, EGFP-Rab7 and EGFP-Rab11a, the corresponding sequences were amplified from mouse brain cDNA and cloned between XhoI and BamHI sites in the pEGFP-C1 vector (a gift from Hilal Lashuel, Swiss Federal Institute of Technology)^93^: Rab5 For: 5’-TCTCGAGCTCAAGCTTTAATGGCTAATCGAGGAGCAACA-3’

Rab5 Rev: 5’-TAGATCCGGTGGATCCTCAGTTACTACAACACTGGCTTCTGG-3’

Rab7 For: 5’-TCTCGAGCTCAAGCTTATGACCTCTAGGAAGAAAGTGTTG-3’

Rab7 Rev: 5’-TAGATCCGGTGGATCCTCAACAACTGCAGCTTTCTG-3’

Rab11a For: 5’-TCTCGAGCTCAAGCTTATGGGAACACGCGACGACGTA-3’

Rab11a Rev: 5’-TAGATCCGGTGGATCCGATGTTCTGACAGCACTGCACCTTT-3’.

T34N and Q75L mutations were introduced in pDEST47-Arl8b^WT^-GFP plasmid (Addgene #67404), using the QuikChange II XL Site-Directed Mutagenesis kit (Agilent). To generate untagged Arl8b constructs, the Arl8b coding sequence was cloned in the pBI-CMV3 bidirectional promoter vector (Clontech). The KLC1-TAP plasmid was a gift from Larry S. B. Goldstein (University of California San Diego). 2xmCh-KIF5C was designed by subcloning 2xmCh from KIF5C(1-560)-2xmChEF(C) (Addgene #61664) in pCDNA3.1. Then, full length KIF5C sequence was amplified from pGFP-KIF5C (a gift from Michelle Peckham, University of Leeds)^94^, and cloned downstream of 2xmCh. To generate the SKIP 1-186 construct, the sequence of the first 186 amino acids was cloned in pcDNA3.1 at the HindIII and BamHI site.

SKIP 1-186 For: 5’ – CTAGCGTTTAAACTTAAGCTTATGGAGCCGGGGGAGGTGAAG – 3’

SKIP 1 -186 Rev: 5’ – CCACACTGGACTAGTGGATCCGACCGAGCTGGGAAGGCGGTC – 3’

### Transfection

Hippocampal neurons were transiently transfected using Lipofectamine 2000 (Thermo Fisher), following the manufacturer instructions. DNA and Lipofectamine were diluted in non-supplemented Neurobasal-A medium (Gibco). For most experiments, neurons grown on 12 mm coverslips in 24-well plates were transfected using 2 µL of Lipofectamine and 0.4 or 0.8 µg of DNA per well. Otherwise, neurons were grown in microfluidic chambers and transfected using 1.2 µL of Lipofectamine and 0.5 µg of DNA per chamber. Medium was changed 1 hour after transfection.

Neuro 2a cells were transiently transfected using Lipofectamine 2000 (Thermo Fisher) following the manufacturer instructions. DNA and Lipofectamine were diluted in DMEM (Gibco). Experiments were performed in 6 well plates, using 8 µL of Lipofectamine and 2 µg of DNA per well.

### Adenovirus transduction

Cre-recombinase adenovirus transduction (University of Iowa, Ad5CMVCre) was performed as described previously^50^. Briefly, hippocampal neurons isolated from Kinesin-1B conditional knockout mice (*KifB^pflox/pflox^)* were treated at 6 DIV with 0, 100, or 400 MOI of adenovirus, corresponding to 0, 5×10^6^, and 2×10^7^ plaque-forming units (PFU), respectively. Neurons were incubated for 2 hours with the virus and then washed twice with Neurobasal-A medium (Gibco) containing 2% B-27 (Gibco), and 0.25% GlutaMAX (Gibco). Neurons were transfected with fluorescently-labeled PrP constructs at 9 DIV and imaged at 11 DIV.

### Adeno-associated virus transduction

PrP^WT^-mCh and PrP^PG14^-mCh were cloned into the pAAV-hSyn-MCS backbone between the BamHI and EcoRI sites (gift from T. Golde, University of Florida).

For: 5’ - CCGCGAGCTCGGATCCATGGCGAACCTTGGCTACTG - 3’

Rev: 5’ - CTTCCTGATCGTGGGATGAGAATTCCTCGAGCAGC - 3’

AAV virus was made using the AAV-DJ Helper Free Packaging System (Cell Biolabs) using published protocols^95^. Hippocampal neurons growing in 24-well plates were treated overnight with 4 µl of the viral stock at DIV7 and imaged at DIV12.

### shRNAs & siRNAs

To knockdown Arl8b from neurons, the following 21-mer shRNA inserts were cloned separately in the pLKO.3G vector (Addgene plasmid #14748): scrambled shRNA (CCTAAGGTTAAGTCGCCCTCG), Arl8b shRNA #1 (CCTCTCGAAATGAACTGCATA), Arl8b shRNA #3 (CGAGGAGTCAATGCAATTGTT). Dynein was knocked down by transfection of neurons with a previously validated DHC1 lentiviral construct (GTGATGCCATACGAGAGAA) and the scrambled construct (GCACACGTATCGACGTATC)^50^. Validation of shRNA constructs for reduction of Arl8b was done by transfecting mouse N2a cells at 40% confluency with pLKO3.G-Arl8b shRNA #1 and #2 either separately or together using Lipofectamine 2000 (Invitrogen). Transfection rates were > 80% and were ascertained by counts of GFP fluorescence on cell bodies. ShRNAs were expressed for 48 or 72h prior to harvesting, lysed in radioimmunoprecipitation assay (RIPA) buffer and evaluated by Western-blot using rabbit anti-Arl8b (Proteintech) and anti-alpha-tubulin as a loading control (Sigma Aldrich). N2a cells were harvested at the same time points for reverse-transcriptase (RT) PCR generation using iScript cDNA Synthesis kit (Bio-Rad), and quantitative PCR (QPCR) was performed using the FastStart Universal SYBR Green Mastermix (Roche) to test for reduced Arl8b mRNA levels.

To knockdown VPS41 from neurons we used the ON-TARGET plus SMART pool siRNA from Dharmacon. VPS41 siRNA #1: CCAAAGGAACAUUAAACGA, VPS41 siRNA #2: GUUUGUACUGGCGGGAAA, VPS41 siRNA #3: GGAGAAGAAUUUCACGAGA and VPS41 siRNA #4: UGACAUAAGUCUUCGCCCA. Non-targeting siRNA #1: UGGUUUACAUGUCGACUAA, was used as a control.

### Live-imaging microscopy

Live imaging was done in soma and axons of hippocampal neurons at 9-15 DIV, usually two days after transfection, except otherwise stated. In some experiments, fluorescently labeled PrP constructs were co-transfected with soluble EGFP or mCh constructs in order to visualize the neuronal morphology and facilitate the determination of axonal polarity. Coverslips (12 mm), were transferred and flipped onto 35 mm glass bottom dishes (MatTek) containing 2 mL of Neurobasal-A medium (Gibco,) containing 2% B-27 (Gibco) and 0.25% GlutaMAX (Gibco) media. Neurons were imaged using a Nikon Ti-E Perfect Focus inverted microscope equipped with a total internal reflection fluorescence microscopy (TIRFM) setup, with an Andor iXon + DU897 EM Camera, and a 100X/1.49 NA oil objective. A 488 nm laser was used for detecting GFP and a 561 nm laser to detect mCh. Lasers were positioned at varying angles for pseudo-TIRFM acquisition. Transfection rates were ∼1%, which enabled imaging of individual neurons. Time-lapse movies of axonal transport had different duration and framerates depending on the type of cargo analyzed. Movies of fast vesicular transport were 15 sec long and collected at a frame rate of 10 frames/sec (10 Hz). Movies of aggregate transport were 5 min (300 sec) long and collected at a frame rate of 1 frame/sec (1 Hz). Movies of LAMP-1-EGFP vesicular transport were 1 min long and collected at a frame rate of 5 frame/sec (5 Hz). For all images and movies acquired, exposure time was set to 100 ms. Pixel size was 0.16 µm. Plates with cultured neurons were maintained at 37°C and 5.5% CO_2_ throughout the total imaging period. All axonal transport movies were taken in a central region of the axon, at least 150 µm away from both the soma and the axon tip.

For green/red axonal cotransport analyses, we performed near-simultaneous two-color imaging. Exposure times were set to 75 ms and red and green images were collected with a 50 ms delay between them using Nikon’s proprietary LAMBDA 10-3 optical filter switch. For axonal transport live imaging, high-resolution imaging of vesicle cotransport was done using a high frame rate: 30 seconds long and at 5 frames/second (5 Hz). Low frame rate cotransport movies were obtained for analyses of aggregate transport: 5 min (300 seconds) long and collected at 1 frame/sec (1 Hz). Time-lapse imaging was performed during 5 min at a frame rate of 5 frames per sec.

### Fixation and Immunofluorescence

Neurons were fixed with 4% paraformaldehyde (PFA, Electron Microscopy Services) containing 4% sucrose (Sigma) for 30 min at 37°C. Cells were washed once in 50 mM Glycine in PBS and 3 more times in PBS. If applicable, cells were permeabilized by incubating the coverslips in 0.1% Triton X-100 (Sigma) in PBS for 8-10 minutes followed by 3 washes with PBS. Coverslips were incubated in blocking solution (10% IgG-Free BSA, 5% donkey serum or goat serum in PBS, Jackson ImmunoResearch) for 30 minutes at room temperature (RT). Primary antibodies were incubated in blocking solution for 1 hour at RT or overnight at 4°C. After 3 washes in PBS, secondary antibodies were incubated in blocking solution for 1 hour at RT or overnight at 4°C. If applicable, neurons were counterstained with 300 nM DAPI (Thermo Fischer) for 5 minutes. After 3 washes in PBS, coverslips were washed once in H_2_O and mounted in ProLong Diamond antifade reagent (Thermo Fisher).

The following antibodies were used for immunofluorescence (IF): mouse anti-KDEL (1:100, Santa Cruz Biotechnology); rabbit anti-GM130 (1:100, Abcam); goat anti-mouse Cathepsin B (1:40, R and D systems); rabbit polyclonal IgG anti-mCh (1:100 Gentex), recombinant Fab anti-PrP (1:200 clone HuM-D13)^96^, rabbit anti-Arl8b (1:200, ProteinTech), and mouse anti-Vps41 (1:100 clone D-12, Santa Cruz Biotechnology). The following antibodies were used for Western blot (WB): rabbit anti-Arl8b (1:200, ProteinTech) and mouse anti-alpha-tubulin (1:500, Sigma).

### Correlative Fluorescence and Scanning Electron Microscopy (SEM)

Hippocampal neurons were plated in microfluidic chambers and co-transfected at 8 DIV with PrP^PG14^-mCh and soluble EGFP. Two days after transfection, the neurons inside entire microfluidic chamber were imaged using live-fluorescence microscopy (Nikon S Plan Fluor ELWD 20X/0.45 [infinity]/0-2 Objective) to map the location of transfected neurons. Shortly thereafter, neurons were initially fixed in ice-cold 2.5% glutaraldehyde in 0.1 M Na cacodylate buffer (pH 7.3) with the addition of 1% trehalose during which the mold was carefully removed to expose fully the cultured cells. After a buffer wash the cells were fixed in 1% osmium tetroxide, washed thoroughly with distilled water and dehydrated in graded ethanol series. Following the 100% ethanol dehydration, the samples are immersed in 100% hexamethyldisilazane (HMDS) for 3 minutes and allowed to dry. Each coverslip was mounted on a stub with carbon tape and samples were sputter coated with iridium at 10μA to a thickness of approximately 5 nm (EMS model 150T S). The coverslips were then examined on a Hitachi S4300 SEM (Hitachi High Technologies America Inc., Pleasanton CA) at 5kV with settings adjusted according to needs.

### Correlative Fluorescence and Serial Sectioning Scanning Fluorescence Electron Microscopy (S3EM)

Hippocampal neurons were plated on 35-mm gridded #1.5 glass bottom dish (Cellvis) pretreated with 50 µg/ml poly-L-lysine (Sigma) in Borate buffer and transfected at 8 DIV with PrP^PG14^-mCh using Lipofectamine 2000 (Thermo Fisher). Two days after transfection, neurons were first imaged using live-fluorescence and phase-contrast microscopy (Nikon S Plan Fluor ELWD 20X/0.45 [infinity]/0-2 Objective) to map the location of transfected neurons on the grid (**Supplementary Fig. 3e**). Then PrP^PG14^-mCh aggregate sites were imaged at high resolution using super-resolution radial fluctuation (SRRF) microscopy^97^ (20 frames/second, 100 frames) with a pseudo-total internal reflection fluorescence microscopy (pTIRFM) setup, with an Andor iXon + DU897 EM Camera, and a 100X/1.49 NA oil objective. Samples were then fixed on the gridded culture dish while imaging on the microscope by carefully mixing an equal volume of 2X EM-fixative (5% glutaraldehyde, 4% paraformaldehyde in 0.1M cacodylate buffer with 3mM CaCl_2_) warmed to 37°C with 2 mL of Neurobasal-A medium (Gibco,) containing 2% B-27 (Gibco) and 0.25% GlutaMAX (Gibco). Samples were immediately removed from the microscope, fixative-media solution was discarded, and cells were left in fresh ice-cold 1X fixative for 90 minutes and rinsed three times with ice cold 0.1M cacodylate buffer with 3mM CaCl_2_. Coverslips were post-fixed and stained with 1.5% reduced osmium for 35 minutes, rinsed five times with MilliQ water, and stained again with 1% aqueous uranyl acetate for 1 hour at room temperature, before serial dehydration with graded solutions of ice-cold ethanol in water. Samples were then fully dehydrated in two 10 minute rinses of anhydrous ethanol at room temperature before infiltration with Durcupan resin. After 3:1, 1:1, 1:3 ethanol-resin two-hour infiltration steps, samples were infiltrated with pure resin for two hours before another change of fresh pure resin and left overnight at room temperature. In the morning, a final change of fresh resin was performed, taking care to let the viscous resin fully drain by inverting the coverslip before adding the final aliquot of fresh resin from the edge of the dish. Resin was added to fill the dish up to 2 mm and infiltrated samples were polymerized for 48 hours at 65°C.

After polymerization, coverslips were dissolved by immersion in concentrated hydrofluoric acid. Correlative light-electron microscopy (CLEM) was achieved using laser branded fiducials in a thinly embedded sample, as previously described^98^. Briefly, regions of interest (ROIs) that included SRRF-imaged PrP^PG14^-mCh axon segments were identified by their grid positions and dark osmium staining. ROIs were marked using the cutting laser of a Zeiss PALM laser cutting microscope to provide orientation and fiducial guides for further trimming and ultramicrotomy. ROIs were carefully identified under a dissecting microscope and excised from the coverslip using a jeweler saw and a scalpel, with careful thought given to the future blockface orientation. The small (1×2×2mm) sample block was glued to a blank resin block such that the ROIs were orthogonal to the cutting plane and approximately 100 µm from the cutting surface. Laser marks on the block face helped to identify the location of the ROI using the ultramicrotome optics. The block was carefully trimmed using a 90° diamond trimming knife (Diatome) so that the block width bounded the fiducial laser marks, and the two sides of the block face were perfectly parallel for serial sectioning when turned 90°.

The ROI was approached using the trimming knife to provide a perfectly smooth block face. When the fiducials marking the location of ROI became challenging to visualize (about 5-15µm from the blockface), a diamond knife with a water boat appropriate for serial sectioning (Histo 45°, Diatome) was installed on the ultramicrotome and 50nm serial sections were collected on a series of plasma cleaned custom-diced silicon chips (University Wafer, Boston, MA) immersed in the knife boat. About 200 serial sections were collected in total over four chips.

The chips were mounted on aluminum stubs using double-sided carbon sticky tape and loaded into a Zeiss Sigma VP scanning electron microscope (SEM). Chip mapping and the imaging of serial sections was facilitated by SmartSEM (Zeiss) and Atlas 5 (FIBICS) software packages. Maps of all of the serial sections on the silicon chips were collected at 500nm/px. Laser marks could be identified in the resin boundary at low magnification, demarcating the ROI in each section. The ROI of the axon segment containing the PrP^PG14^-mCh aggregates was identified and captured across 40 serial section scanning electron micrographs. ROIs were imaged at 2nm/px using an electron backscatter detector (Gatan). In high-vacuum mode, a 3keV beam in high-current mode with a 30 µm aperture at a working distance of 9 mm was found to produce signal from which we could resolve membrane and organelle (eg., microtubule) ultrastructure. Sections were aligned using Photoshop (Adobe) and elastic alignment functions^99^ embedded in Fiji (NIH).

The orthogonal projection (eg., x-z) of the S3EM data was used to correlate the SRRF-processed data and produce overlay (**Fig. 3e and Supplementary Fig. 3e**). Sections were sampled from throughout the aggregate and non-aggregate sites with reference to the correlated volume.

### Hippocampal Neuron Live Assays with Various Drug Treatments and Fluorescent Dyes

To retain PrP^PG14^ in the ER, Brefeldin-A (BFA) (BioLegend) was applied at 5 µg/mL at the same time as PrP^WT^-mCh or PrP^PG14^-mCh transfections and incubated for 2-6 hours. Neurons are washed 2-3 times with Neurobasal-A medium (Gibco) prior to fixation.

For characterization of PrP^PG14^ at the cell surface of the soma or axons, neurons grown in 24-well plates, were transfected with a PrP^PG14^-mCh-BBS construct for 1 hour. Neurons were treated with AlexaFluor647-labeled-α-Bungarotoxin (BTX-a647, Molecular Probes) by applying it to the media and as described previously^66^. Briefly, we pretreated neurons for 1 hour with 100 µM tubocurarine hydrochloride (Sigma), to eliminate non-specific binding to other receptors^66^. Neurons were then incubated with 7µg/mL BTX- a647 in the media, during indicated times and at indicated temperatures. To inhibit clathrin mediated-endocytosis, 80 µM Dynasore (Sigma) was added 30 minutes prior to BTX-a647 treatment and kept during the labeling. DMSO was used as a control. Internalized PrP^PG14^ particles were defined as PrP^PG14^-mCh-BBS particles that were both positively labeled with BTX-a647 and that were actively transporting within axons. Quantification of internalization in the presence or absence of Dynasore was performed by measuring the proportion of BTX-a647 positive particles that were actively transporting per movie.

PrP^PG14^ surface expression pattern in axons was analyzed by incubating neurons with BTX-a647 during 10 minutes at 4°C before fixation. LysoTracker Green (Molecular Probes) was incubated at 50 nM, 1-2 hours before live-imaging. Magic Red from the Cathepsin B Assay (ImmunoChemistry Technologies), was incubated 1-2 hours before live-imaging. 260X DMSO stock of Magic Red dye was diluted in sterile diH_2_O and 10 µL was applied onto neurons in each well containing 250 µl of Neurobasal-A medium (Gibco). Bafilomycin A1 (Sigma) was incubated at 10 nM during 6hours. Longer incubation times affected neuronal survival. DMSO was used as a control.

For characterization of PrP^PG14^ cleavage at the cell surface, hippocampal neurons were transfected with PrP^PG14^-mCh-TCS or PrP^PG14^-mCh. Neurons were incubated with or without 5 units/ml of Thrombin protease (Sigma) 2 hours after transfection. This treatment was maintained for 48 hours. We did not observe any effect of 5 units/ml Thrombin treatment on neuronal survival.

For characterization of (in)dependence between direct and indirect ARESTA, hippocampal neurons were transfected with PrP^PG14^-mCh-TCS and Arl8b^WT^-GFP. Neurons were incubated with or without 5 units/ml of Thrombin protease (Sigma) 24 hours after transfection. This treatment was maintained for 24 hours. We did not observe any effect of 5 units/ml Thrombin treatment on neuronal survival.

### Photoactivation of PrP^PG14^-PAmCh

Neurons were cotransfected with EGFP-Rab7 and PrP^PG14^-PAmCh. Two days after transfection, PrP^PG14^-PAmCh was photoactivated using a 405 nm laser burst pointed at the cell body (40% laser power, 3 sec exposure), and neurons were imaged using live-fluorescence microscopy. The fate of the photoactivated molecules and their delivery into axons was followed for 60 minutes, by taking 5×5 stitching images every 5 minutes.

### Proteinase-K Resistance Assay

N2a cells were transfected with PrP^WT^-mCh or PrP^PG14^-mCh (pcDNA3.1 vector), and lysed 48 hours after transfection in PBS containing 0.5% Nonidet P-40 (Sigma), 0.5% sodium deoxycholate (Sigma), 0.2% Sarkosyl (Sigma), and 0.5% Triton X-100. Proteins in 500 µg of lysate were digested using 0, 0.05, 0.1, 0.5, 1 or 5 µg/ml of Proteinase K (PK) (Thermo Fisher), during 30 minutes at 37°C. Reaction was blocked by adding 0.8 mg/ml of freshly prepared PMSF (Sigma). The equivalent of 20 µg of each sample was analyzed by immunoblotting using an anti-PrP specific antibody (D13) (gift of D. Burton, Scripps Research). Samples corresponding to 1 and 5 µg/ml of PK were also precipitated using trichloroacetic acid (TCA) (Sigma) to concentrate the proteins. The equivalent of 120 µg of each sample was analyzed by immunoblotting.

### SDS-Page and Immunoblotting

Neuronal lysates were resuspended in Laemmli buffer and resolved by SDS-PAGE in 8– 12% polyacrylamide gels. Proteins were electrotransferred onto PVDF membranes. Membranes were blocked for 30 minutes at RT in TBS containing 0.1% Tween 20 and 5% dry milk. They were incubated overnight at 4°C with primary antibodies diluted in TBS-Tween-milk. After six washes in TBS-Tween, the membranes were further incubated for 1h at RT with fluorescent secondary antibodies (Li-Cor) diluted in TBS-Tween-milk. After 6 more washes in TBS-Tween, the membranes were imaged using Li-Cor Odissey imaging system using recommended antibodies.

### Calcium Imaging

Hippocampal cultures from WT (C57BL6) or *Kif5c^-/-^* mice were plated directly on #1.5 glass-bottomed dishes (MatTek), at the density of 8-15 neurons per 420 µm^2^ field of view. Cultures were loaded with 2 µM Fluo-4 AM (Thermo Fisher) in Neurobasal-A media supplemented with B27 without phenol red for 30 minutes at 37°C in a 5.5% CO_2_ atmosphere, as recommended by Invitrogen protocol. Cultures were washed 3 times and incubated for another 30 minutes in the B-27 supplemented Neurobasal-A without phenol red prior to imaging to allow complete de-esterification of intracellular AM esters. Prior and during imaging, calcium imaging buffer (140 mM NaCl, 5 mM KCl, 2 mM CaCl_2_, 0.8 mM MgCl_2_, 10 mM glucose, 10 mM HEPES [pH 7.4]) was applied.

Neurons were imaged using a Nikon S Plan Fluor ELWD 20X/0.45 [infinity]/0-2 Objective in a temperature-controlled enclosure set to 37°C. Epifluorescence excitation was provided by a metal-halide-doped mercury arc lamp (Nikon Intensilight). High temporal-resolution images (5 frames/second) were taken for 1 minute with EM-CCD camera (Andor iXon3 897). Fluo-4 AM fluorescence were collected using Chroma filters (ET-GFP [ET470/40x T495lpxr, ET525/50m]. Depolarization was induced by applying KCl to a final concentration of 30 mM at the 30-second time point during acquisition. Still images of PrP^WT^-mCh or PrP^PG14^-mCh of the same region were collected using ET-DsRed [ET545/30x, T570lp, ET620/60m] Chroma filter.

### Cell Death Assay and Imaging

ReadyProbes® Cell Viability Imaging Kit (Blue/Green) (Thermo Fisher) was used to quantify cell death. In brief, two drops of NucBlue® Live reagent (Hoechst 33342) and two drops of NucGreen® Dead reagent were added onto 10-14 DIV hippocampal neurons grown on the coverslips containing 1 ml of Neurobasal-A medium (NBA) (Gibco), supplemented with 2% B-27 (Gibco) and 0.25% GlutaMAX (Gibco). Neurons were incubated at 37°C for 30 minutes prior to fixation with 4% PFA (Electron Microscopy Services) containing 4% sucrose (Sigma) for 30 min at 37°C. Cells were washed once in 50 mM Glycine in PBS and 3 more times in PBS before mounted in ProLong Diamond antifade reagent (Thermo Fisher).

Fixed neurons were imaged using a Nikon S Plan Fluor ELWD 20X/0.45 [infinity]/0-2 Objective. Epifluorescence excitation was provided by a metal-halide-doped mercury arc lamp (Nikon Intensilight). Blue and Green nuclei staining were collected using UV-2E/C [AT350/50x, T400lp, ET460/50m] and ET-GFP [ET470/40x, T495lpxr, ET525/50m] Chroma filters respectively. PrP^WT^-mCh or PrP^PG14^-mCh fluorescence were collected using ET-DsRed [ET545/30x, T570lp, ET620/60m] Chroma filter.

## QUANTIFICATION AND STATISTICAL ANALYSIS

### Neuronal Soma Image Analysis

All images of neuronal soma were processed in ImageJ (National Institutes of Health) ^100^. Rolling-ball background fluorescence correction was applied to all images. 3D Deconvolution was applied using synthetic Gaussian point spread function (PSF) and Richardson-Lucy (RL) algorithms in DeconvolutionLab2^101^. All images and line scan fluorescence intensity graphs were displayed as maximum intensity projection.

### Analysis of Axonal Transport Dynamics from Time-lapse Movies

Axonal transport analysis was performed using the custom-made and previously validated *KymoAnalyzer* ImageJ package of macros ^70^. *KymoAnalyzer* is freely available for download from our lab website (http://www.encalada.scripps.edu/kymoanalyzer), and a detailed description are available here and in ^70^. Briefly, kymographs were generated from time-lapse movies. Particle trajectories were manually assigned from the kymograph images. Track and segment-related parameters were automatically calculated by KymoAnalyzer.

### Colocalization and Cotransport Quantifications

We determined colocalization by merging line-scan intensity profiles of different channels in the same graph and quantifying the average number of peaks that overlapped between different channels. For cotransport analyses, individual and merged color kymographs were generated and quantification of the average number of tracks that overlapped between each pair of vesicular cargoes was analyzed. Vesicle tracks were sorted in anterograde, retrograde and stationary populations, or as indicated for individually analyzed parameters. Aggregate tracks were sorted in mobile and stationary populations, or as indicated for individually analyzed parameters.

### Quantification of Particle Densities in Axons

PrP^PG14^ aggregate densities were quantitated from still images of axons transfected with fluorescently-labeled PrP constructs, as follows: fluorescence background was subtracted by quantification of the maximal gray intensity of five small PrP^PG14^ vesicles (as define by visual comparison of size to those vesicles that actively move in live movies), quantified for each image analyzed. The calculated intensity average values were subtracted from each image using the Brightness and Contrast tool of ImageJ. The particles still present in the image after thresholding were considered as aggregates. Axon length was measured using the polyline tool of ImageJ. Aggregate density was expressed as the average number of aggregates per 100 µm of axon length.

For Lysotracker-positive vesicle densities, we measured the axon length using the polyline tool of ImageJ and quantified all the particles labelled by Lysotracker-Green along this line. Densities were plotted per 100 µm of axon length.

### ERSE-mCh and Vps41 Fluorescence Quantification

Neurons were cotransfected with the pEGFP-C1, moPrP.Xho PrP^WT^-EGFP or moPrP.Xho PrP^PG14^-EGFP and the transcriptional ER stress reporter ERSE-mCh (a gift from Larse Plate, Vanderbilt University) ^64^ Increases in the activity of ER stress pathways activate ERSE and soluble mCh is synthesized.

We quantified total fluorescence intensity (gray values, arbitrary unit, a.u.) of ERSE-mCh and Vps41 using ImageJ, using circular ROIs of 30 µm in diameter centered around the soma. For axonal total fluorescence intensity, polyline tool was used to draw ROIs that cover the entire axonal length.

### Calcium Imaging Analysis

Regions of interest (ROIs) were marked around soma of neurons expressing PrP^WT^-mCh or PrP^PG14^-mCh, and we measured average fluorescence for each ROI at a given timepoint. The average Fluo-4 AM fluorescence overtime was collected using an ImageJ Plugin Time Series Analyzer V3 and normalized to *Δ*F/F by subtracting fluorescence intensity of each time point (F) to an average fluorescence prior to addition of KCl (F_0_) and divided by F. Rolling-ball background fluorescence correction was applied to all images prior to fluorescence intensity analysis.

### Cell Death Quantitation

All image processing was done in ImageJ. The total number of NucBlue® Live positive nuclei (blue channel) were determined as follows: background fluorescence was corrected on each image, by applying a Gaussian blur filter with a radius of 12 to a duplicate of each image and subtracting this filtered image from the original. An automatic threshold was applied to the resulting images using the Li algorithm followed by a watershed filter, both in ImageJ. Total nuclei were counted using a minimum size threshold of 15 and circularity of 0.10. The total amount of NucGreen® Dead positive nuclei (green channel) were determined using the same method, now using a Gaussian blur filter with a radius of 10, thresholding using the Yen algorithm available in ImageJ, and nuclei determination using a minimum size threshold of 15 and circularity of 0.10.

### Generation of Graphs and Figures

Graphs of average values were generated using Microsoft Excel. Cumulative frequency graphs were generated using the CDF function in MATLAB (Mathworks). Box plots were generated using BoxPlotR, an application in the shiny package from RStudio (http://shiny.chemgrid.org/boxplotr/). Figures were drawn in Adobe Illustrator. Neuron cartoon (graphical abstract) was downloaded from Biorender (https://biorender.com/).

### Statistical Analyses

All the transport parameters measured in this study were first tested for normality using the MATLAB Lilliefors test. For parameters following a normal distribution, we performed a Student’s t-test. For non-normal parameters, we performed a permutation t test (rndttest function in MATLAB). For all parameters, differences in medians were also checked using the Wilcoxon rank-sum test (rnk function in MATLAB). The non-parametric Kolmogorov-Smirnov test was used to evaluate the equality of two sample distributions (in cumulative distribution functions). Most of the parameters are presented as mean ± SEM. Multiple comparison corrections were used where appropriate. Details about each parameter, including definition of center, value and definition of n, statistical test used and p values, can be found in specific figures and/or in figure legends.

## ACKNOWLEDGEMENTS

We thank Malcom Wood at the Scripps Research EM core for performing the SEM experiments; David Harris (Boston U.), Lars Plate (Vanderbilt U.), Larry S. B. Goldstein (U. California San Diego), and Vladislav Verkhusha (Albert Einstein College of Medicine) for sharing reagents; Susan Ackerman, Jeff Kelly, Justine Lebeau, Malene Hansen, Michael Petrascheck, Luke Wiseman, Federico Zampa, Giordano Lippi, George Campbell, Kayal Madhivanan, and all members of the Encalada Lab for advice and valuable discussions and comments. This work was supported, in part, by a NIH/NIA R01AG049483 grant; by the Glenn Foundation for Medical Research Award for Research in Biological Mechanisms of Aging; by a New Scholar in Aging Award from the Lawrence Ellison Foundation; and by a Baxter Family Foundation award to S.E.E. R.C. was supported by a fellowship from the George E. Hewitt Foundation for Medical Research. T.C. was supported by a Royal Thai Government Scholarship from the Development and Promotion of Science and Technology Talents Project (DPST). U.M., S.W.N., and L.R.A. are supported by the Waitt Foundation, Core Grant application NCI CCSG (CA014195), NSF NeuroNex Award No. 2014862, and NIH/NIDCD R21 award DC018237.

## AUTHOR CONTRIBUTIONS

S.E.E., and R.C. conceptualized the research. R.C., A.V., T.C, S.W.N, L.R.A. performed the research. A.V., T.C., S.W.N, L.R.A. validated the research. R.C., A.V., T.C., S.W.N, L.R.A., U.M., S.E.E., analyzed the data. R.C., S.E.E wrote the original draft. S.E.E., T.C., A.V., U.M. reviewed, edited manuscript.

## Additional Information

Supplementary Movie 1. Degradation of PrP^PG14^-mCh in lysosomes in the soma.

Movie of a neuron co-expressing LAMP1-EGFP (green) and PrP^PG14^-mCh (red) showing PrP^PG14^-mCh puncta disappearing. (**Related to Supplementary Fig. 1g**).

Supplementary Movie 2. Axonal transport of PrP^WT^-mCh vesicles.

Inverted contrast movie showing axonal transport of PrP^WT^-mCh vesicles in an axon of a cultured hippocampal neuron. (**Related to Fig. 4 and Supplementary Fig. 4a**).

Supplementary Movie 3. Axonal transport of PrP^PG14^-mCh vesicles and aggregates.

Inverted contrast movie showing axonal transport of PrP^PG14^-mCh vesicles and aggregates in an axon of a cultured hippocampal neuron. (**Related to Fig. 4 and Supplementary Fig. 4a**).

**Supplementary Figure 1.**
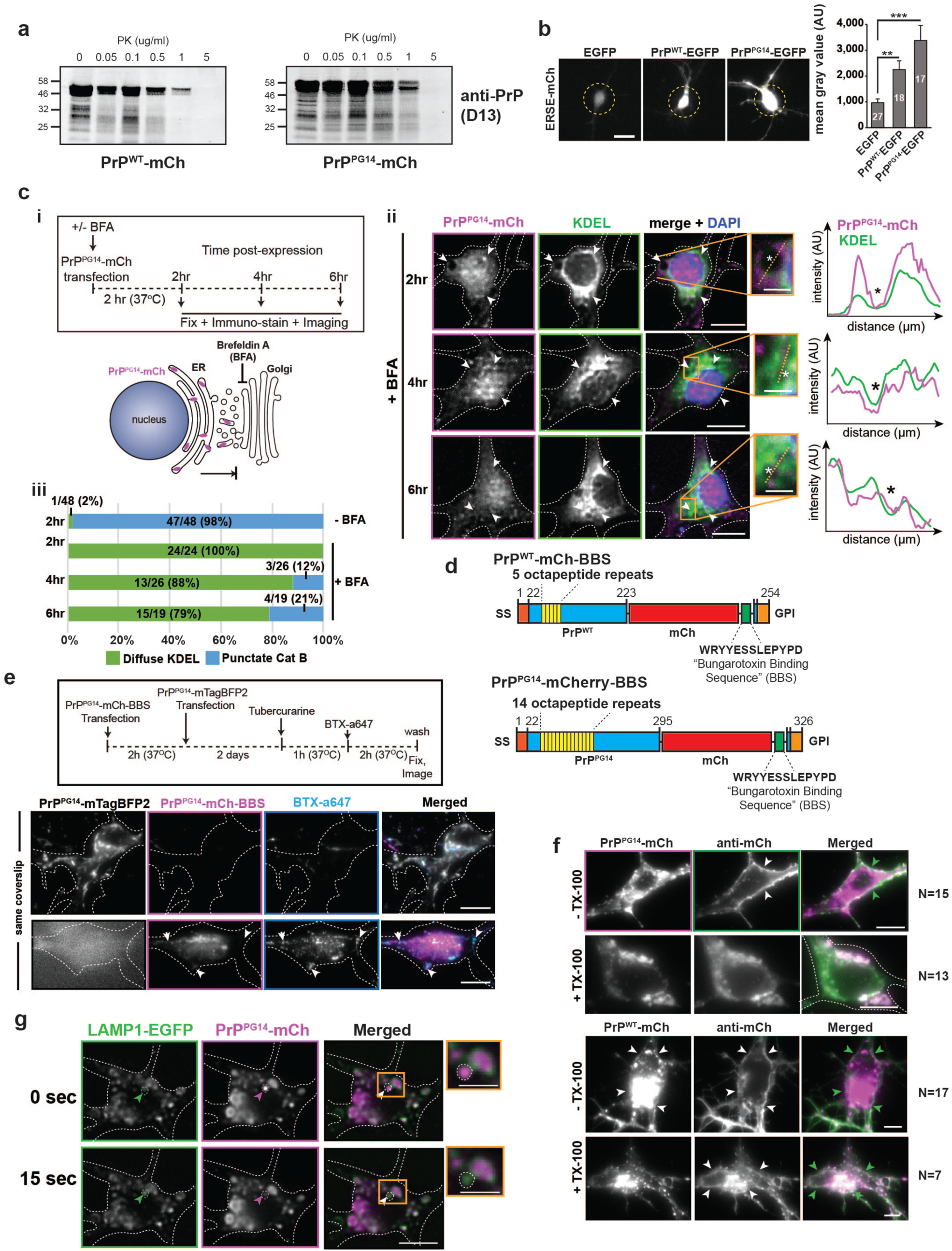
**PrP^PG14^ clearance Post-ER stress and cell surface export in neuronal soma. Related to Fig. 1. a**, Representative Western blots showing of partial Proteinase K (PK) resistance by PrP^PG14^ aggregates from N2a cell lysates (N_replicates_ = 3). **b**, Representative images and quantitation of neurons co-transfected with ERSE-mCh and EGFP (control), PrP^WT^-EGFP, or PrP^PG14^-EGFP. Dotted lines indicate 30 μm diameter circles used to measure total gray intensity values. Scale bar = 20 μm. Values are shown as mean ± SEM. ***p<0.001, **p<0.01, Student’s t-test. N_cells_ are shown inside bars. **c**, **(i)** Experimental outline (top) and diagram (bottom) of BFA assay. **(ii)** Representative images of soma of neurons expressing PrP^PG14^-mTagBFP2 and stained with antibodies against ER (KDEL) or with DAPI nuclear marker at indicated time points in the presence of BFA. Arrowheads and asterisks point to colocalization or non-colocalization events, respectively, also shown in enlarged insets and linescan intensity profiles. Scale bars = 10 μm. Scale bars of insets = 250 nm. **(iii)** Quantitation of PrP^PG14^-mCh colocalization with KDEL or CatB from Fig. 1b. Numbers inside/above bars are total number and percentage of observed neurons. **d**, Schematic of PrP^WT^-mCh-BBS and PrP^PG14^-mCh-BBS constructs. SS: signal sequence; GPI: GPI anchor. **e**, Experimental outline of BBS internalization assay (top). Representative images of two neurons taken from the same coverslip, sequentially transfected with PrP^PG14^-mCh-BBS and PrP^PG14^-mTagBFP2 constructs. Scale bars = 10 μm. **f**, Immunofluorescence images of axons expressing PrP^PG14^-mCh or PrP^WT^-mCh in the presence or absence of detergent (TX-100). Scale bars = 10 μm. **g**, Images of a neuron co-expressing LAMP1-EGFP and PrP^PG14^-mCh at two time points. Scale bar = 10 μm. Arrowheads indicate a PrP^PG14^-mCh disappearing vesicle (enlarged in inset). Scale bars in insets = 5 μm.

**Supplementary Figure 2.**
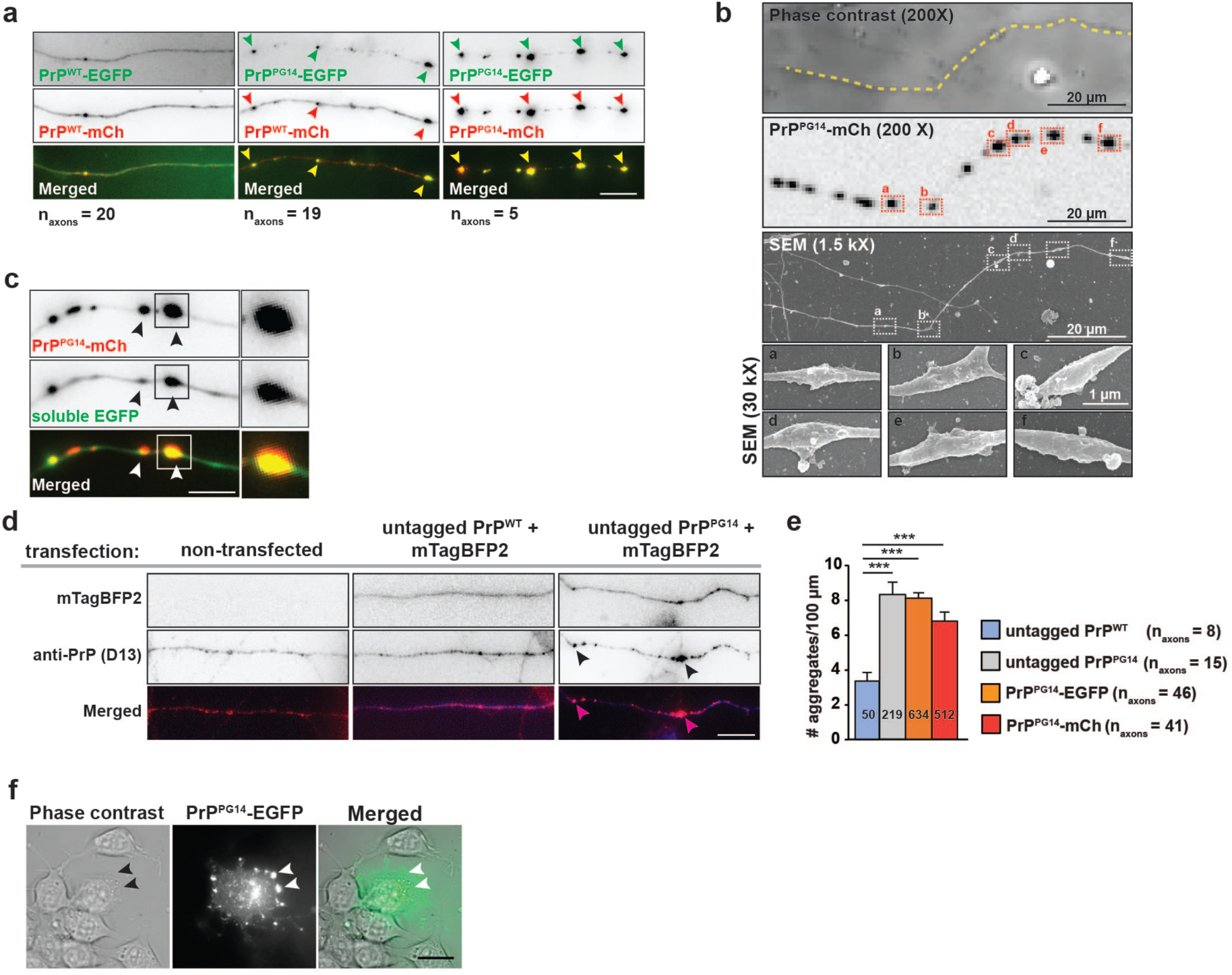
**PrP^PG14^ forms aggregates inside axons. Related to Fig. 2. a**, **I**mages of axons from neurons co-transfected with PrP^WT^-EGFP plus PrP^WT-^mCh (arrowheads point to accumulated PrP^WT-^mCh), or PrP^PG14^-EGFP and PrP^PG14^-mCh (arrowheads point to aggregating PrP^PG14^-EGFP). Scale bar = 10 μm. **b**, Correlative phase contrast (top), fluorescence (middle), and SEM (bottom) images of PrP^PG14^-mCh aggregate swellings. Insets (**a-f**) show enlargements of SEM boxes. N_swellings_ = 20. **c**, Images of axons co-expressing of PrP^PG14^-mCh and soluble EGFP (left panels). Scale bar = 5 μm. Insets of enlargements (right panels). Scale bar in insets = 2.5 μm. Arrowheads point to aggregates. N_axons_ = 39. **d**, Images of non-transfected axons, and those co-expressing untagged PrP^WT^ or PrP^PG14^ with soluble mTagBFP2. Arrowheads point to aggregates. Scale bar = 10 μm. **e**, Quantification of aggregate densities from (**d**). All values are shown as mean ± SEM. ***p<0.001, Kruskal-Wallis test. N_aggregates_ are shown inside bars. **f**, Phase contrast and fluorescence images of a differentiated neuroblastoma (Neuro2a) cell expressing PrP^PG14^-EGFP. Scale bar = 20 μm.

**Supplementary Figure 3.**
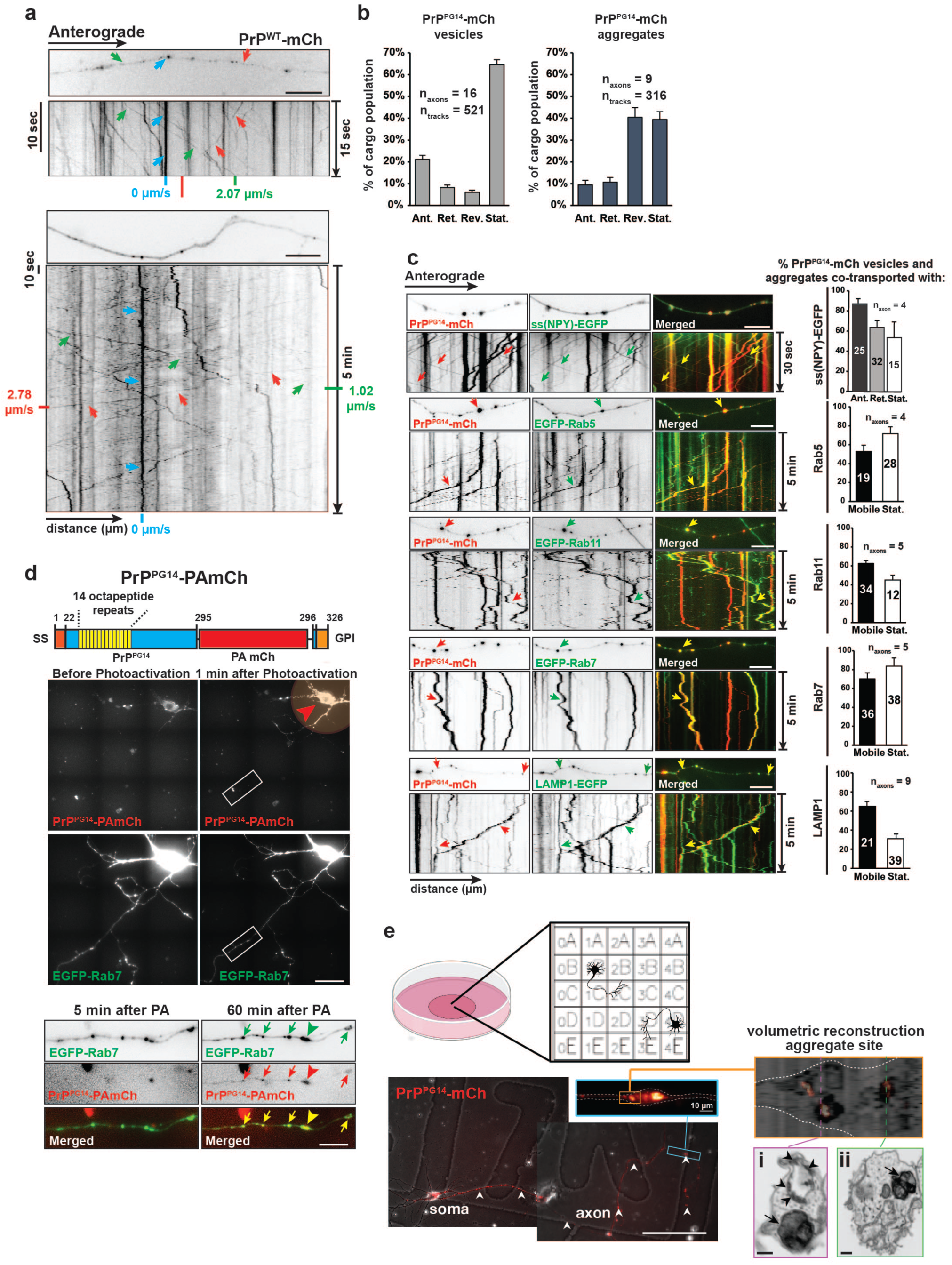
**Post-Golgi PrP^PG14^ vesicles sort to axonal endo-lysosomal compartments. Related to Fig. 3. a**, First frames (top panels of each set) of time-lapse movies of axons expressing PrP^WT^-mCh, and corresponding kymographs (bottom panels of each set) at two time-scales. Arrows point to vesicles moving in the anterograde (green) or retrograde (red) directions, or that are stationary (blue). Scale bars = 10 μm. **b**, Population breakdown of PrP^PG14^-mCh vesicles (left) and aggregates (right). All values are shown as mean ± SEM. **c**, Left panels: Representative first-frame images and kymographs of time-lapse movies of axons co-expressing PrP^PG14^-mCh and various compartment markers: ss(NPY)-EGFP (Golgi-derived secretory), EGFP-Rab5 (early endosomes, EGFP-Rab11 (recycling endosomes), EGFP-Rab7 (LEs), or LAMP1-EGFP (endo-lysosomes). Arrows point to cotransport. Right panels: Quantitation of cotransport between PrP^PG14^-mCh vesicles or aggregates and each of the respective markers. Ant. = anterograde, Ret. = retrograde, Stat. = stationary. All values are shown as mean ± SEM. N_vesicles or aggregates_ are shown inside bars. Scale bars = 10 μm. **d**, Top: schematic of PrP^PG14^-PAmCh construct. SS: signal sequence; GPI: GPI anchor. Middle: representative images of a hippocampal neuron co-transfected with PrP^PG14^-PAmCh and EGFP-Rab7 before (left panels) and after (right panels) photoactivation (pink circle, arrowhead) of the cell body with a 405 nm laser. Inset shows axonal region enlarged in bottom panels. Scale bar of middle panels = 50 μm. Bottom panels: enlargements of inset in middle panels showing axonal PrP^PG14^-PAmCh and EGFP-Rab7 at indicated time points post-photoactivation. Arrows indicate colocalization. Arrowhead points to larger aggregate. N_neurons_ = 3. Scale bar of bottom panels = 10 μm. **e**, Top panels: Schematic diagram of experimental setup showing neurons grown on gridded coverslips. Bottom panels: correlative fluorescence and SEM of neurons expressing PrP^PG14^-mCh. White arrowheads point to axon. Scale bar = 100 μm. Top blue inset shows enlargement of blue rectangle. Right orange inset shows volumetric reconstruction. Magenta and green rectangles show single cross sections at two positions (**i,ii**) of the volumetric reconstruction. Arrowheads point to intralumenal vesicles and arrows point to aggregate structures inside membranous organelles. Scale bar of volumetric reconstruction and insets = 500 nm.

**Supplementary Figure 4.**
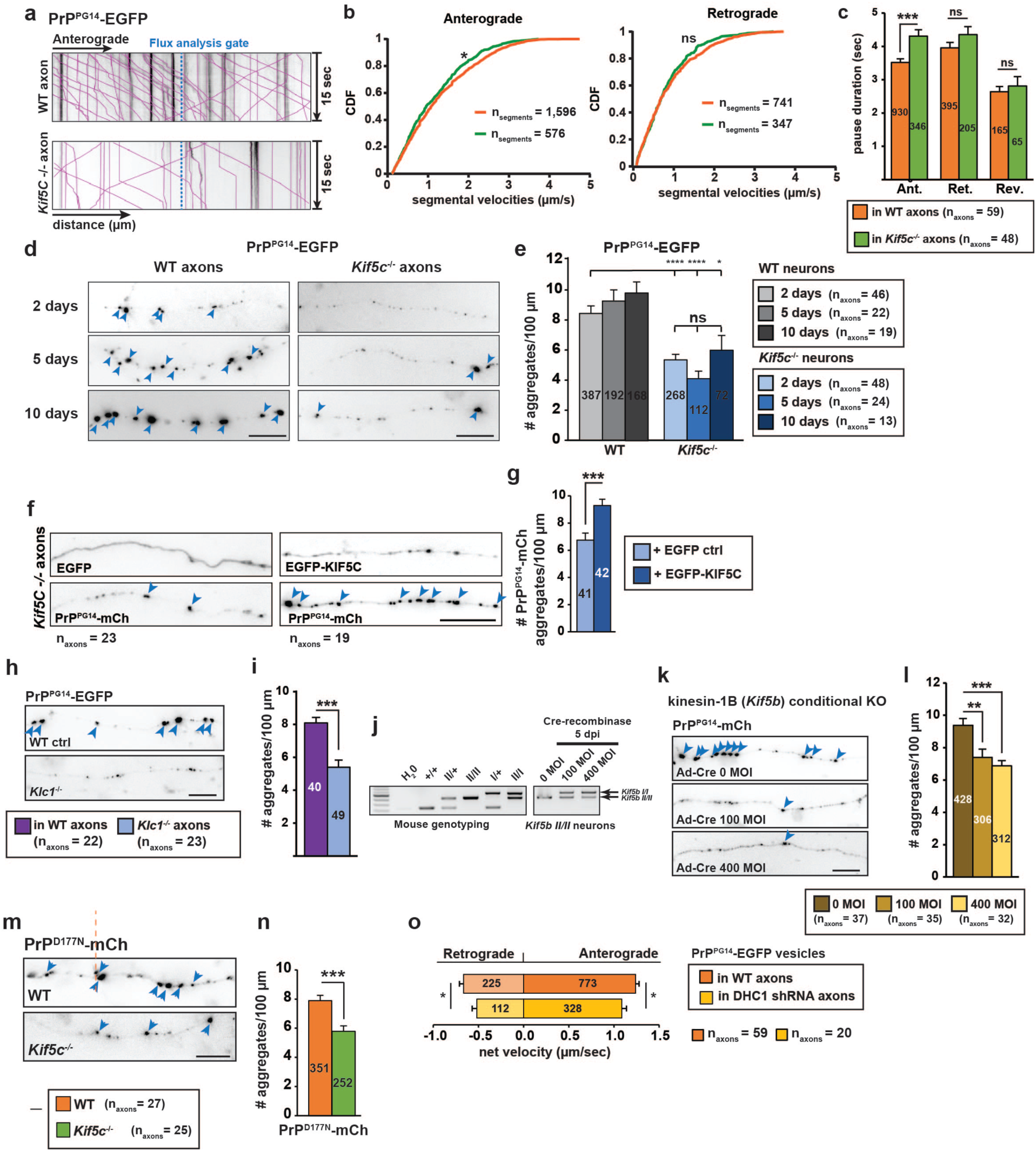
**Kinesin-1- and DHC1-dependent PrP^PG14^ transport and aggregation in axons. Related to Fig. 4. a**, Representative kymographs showing position of gate (dotted blue line) used in flux analyses. **b**, Cumulative distribution frequency (CDF) of anterograde (top) and retrograde (bottom) segmental PrP^PG14^-EGFP vesicle velocities. *p<0.05, Kolmogorov-Smirnov test. **c**, Pause duration of anterograde, retrograde or reversing PrP^PG14^-EGFP vesicles. N_vesicles_ are shown inside bars. **d**, Images of axons from WT (left) and *Kif5c*^-/-^ (right) neurons expressing PrP^PG14^-EGFP at various time points post-transfection. Arrowheads point to aggregates. Scale bars = 10 μm. **e**, Quantitation of aggregate densities from (**d**). **f**, Images of *Kif5c*^-/-^ hippocampal neurons co-expressing soluble EGFP or EGFP-KIF5C and PrP^PG14^-mCh. Scale bar = 10 μm. **g**, Quantitation of aggregate densities from (**f**). N_aggregates_ are shown inside bars. N_aggregates_ are shown inside bars. **h**, Images of axons from WT (top) and *Klc1*^-/-^ (bottom) neurons expressing PrP^PG14^-EGFP. Arrowheads point to aggregates. Scale bar = 10 μm. **I**, Quantitation of aggregate densities from (**h**). N_aggregates_ are shown inside bars. **j**, DNA gel showing genotyping bands for Kinesin-1B hippocampal neuron extracts treated with 0, 100, or 400 multiplicity of infection (MOI) units of cre-recombinase adenovirus (AVV-cre) at 5 days post infection (dpi). The kineins-1B (*Kif5B)* II product is excised upon cre-treatment to convert to a kinesin-1B I/+ product. MOI = Multiplicity of infection. **k**, Images of axons from *Kif5B* conditional knockout neurons expressing PrP^PG14^-mCh at indicated AAV-cre units. Arrowheads point to aggregates. Scale bar = 10 μm. **l**, Quantitation of aggregate densities from (**k**). N_aggregates_ are shown inside bars. **m**, Representative images of PrP^D177N^-mCh aggregates in WT and *Kif5C^-/-^* axons. Scale bar = 10 μm. **n**, Quantitation of aggregate densities from (**m**). N_aggregates_ are shown inside bars. **o**, Average net axonal velocities of PrP^PG14^-EGFP vesicles in WT and DHC1 shRNA neurons. N_tracks_ are shown inside bars. Values in (**c, e, g, i, l, n, o**) are shown as mean ± SEM. *p<0.05, **p<0.01, ***p<0.001, ****p<0.0001, Student’s t-test and Kruskal-Wallis test.

**Supplementary Figure 5.**
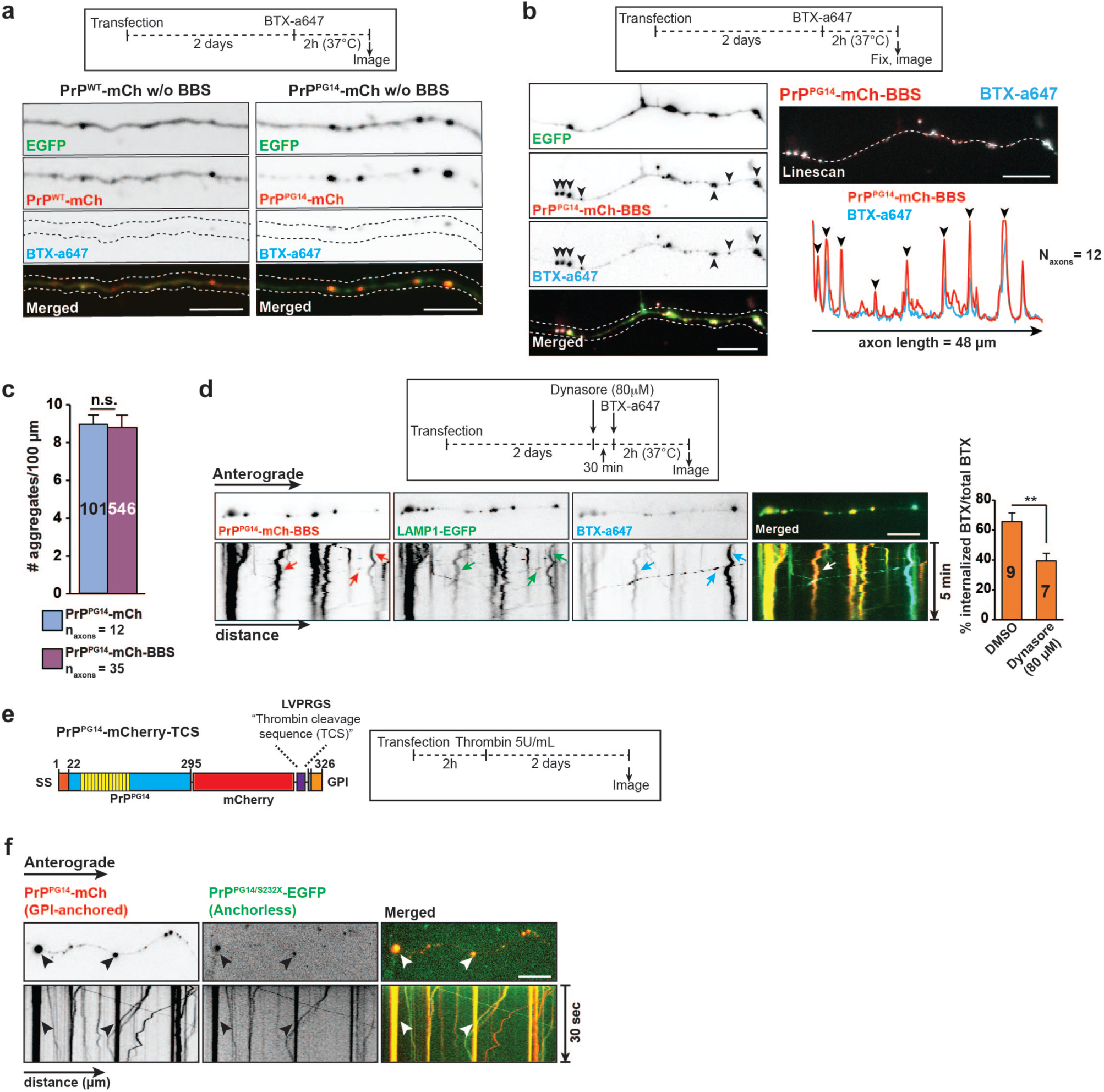
**PrP^PG14^ transiently traffics to the cell surface before intra-axonal aggregation. Related to Fig. 5. a**, Top: Experimental timeline of BTX-a647 labeling assay. Bottom: Representative images of neurons expressing PrP^WT^-mCh or PrP^PG14^-mCh, both without BBS sequence, after treatment with BTX-a647. Scale bars = 10 μm. **b**, Top: Experimental outline of BBS internalization and BTX-a647 labeling assay. Bottom: Representative images and corresponding line scan intensity profile of axons of neurons expressing soluble EGFP and PrP^PG14^-mCh-BBS after BTX-a647 treatment. Arrowheads point to aggregates. Scale bars = 10 μm. **c**, Quantitation of aggregate densities from (**a, b**). N_aggregates_ are shown inside bars. **d**, Top: Experimental outline of Dynasore endocytosis assay. Bottom left : Representative first-frame images of time-lapse movie, and kymographs of axons of neurons co-expressing PrP^PG14^- mCherry-BBS, LAMP1-EGFP and treated with BTX-a647. Arrows point to cotransporting tracks. Bottom right: quantitation of normalized percent internalized BTX-a647 signal. N_axons_ shown inside bars. Values are shown as mean ± SEM. **p<0.01, Student’s t-test. Scale bar = 10 μm. **e**, Left panel: schematic of PrP^PG14^-mCh-TCS construct. SS: signal sequence; GPI: GPI anchor. Right panel: Experimental outline of thrombin assay. **f**, Representative first-frame images of time-lapse movie, and kymographs of axons from neurons co-expressing GPI-anchored PrP^PG14^-mCh and anchorless PrP^PG14/S232X^-EGFP. Arrowheads point to cotransport. N_axons_ = 7. Scale bar = 10 μm.

**Supplementary Figure 6.**
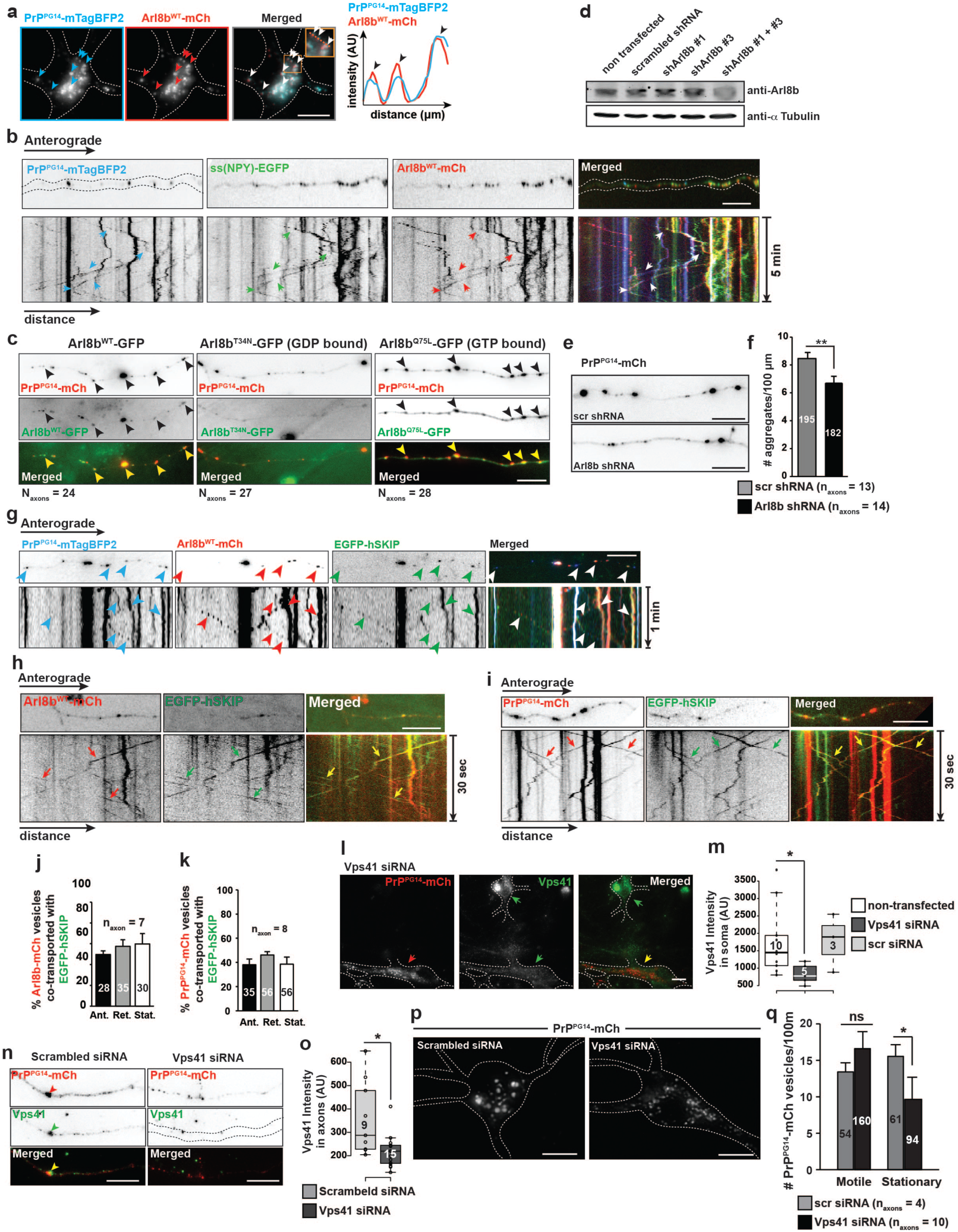
**Arl8b recruits the HOPS complex to PrP^PG14^ vesicles to promote fusion, and aggregation. Related to Fig. 6. a**, Left panels: representative images of hippocampal soma co-expressing PrP^PG14^-mTagBFP and Arl8b^WT^-mCh. Arrowheads point to colocalization. Right pane: line scan intensity profiles of region indicated by dotted orange line inside inset (right). Scale bar = 10 μm. **b**, Representative first-frame images of time-lapse movies, and corresponding kymographs of axons from neurons co-expressing PrP^PG14^-mTagBFP2, ss(NYP)-EGFP and Arl8b^WT^-mCh. Arrows point to cotransport. N_axons_ = 4. Scale bar = 10 μm. **c**, Representative images of neurons co-expressing PrP^PG14^-mCh and Arl8b^WT^-GFP, Arl8b^T34N^-GFP, or Arl8b^Q75L^-GFP. Arrowheads point to colocalization. Scale bar = 10 μm. **d**, Representative Western blot of N2a cell lysates treated with indicated shRNAs. N_replicates_ = 3. **e**, Representative images of axons from neurons expressing PrP^PG14^-mCh treated with scrambled or Arl8b shRNAs. Scale bars = 10 μm. **f**, Quantitation of aggregate densities from (**e**). N_aggregates_ are shown inside bars. **g**, Representative first-frame images of time-lapse movies, and corresponding kymographs of axons from neurons co-expressing PrP^PG14^-mTagBFP2, Arl8b^WT^-mCh and EGFP-hSKIP. Arrowheads point to cotransport. N_axons_ = 15. Scale bar = 10 μm. **h**, Representative first-frame images of time-lapse movies, and corresponding kymographs of axons from neurons co-expressing Arl8b^WT^-mCh and EGFP-hSKIP. Arrows point to cotransport. Scale bar = 10 μm. **i**, Representative first-frame images of time-lapse movies, and corresponding kymographs of axons from neurons co-expressing PrP^PG14^-mCh and EGFP-hSKIP. Arrows point to cotransport. Scale bar = 10 μm. **j**, Quantitation of percentage of cotransporting Arl8b^WT^-mCh and EGFP-hSKIP vesicles from (**H**). N_vesicles_ are shown inside bars. **k**, Quantitation of percentage of cotransporting PrP^PG14^-mCh and EGFP-hSKIP vesicles from (**i**). N_vesicles_ are shown inside bars. **l**, Immunofluorescence images of the soma of neurons expressing PrP^PG14^-mCh, transfected with Vps41 siRNA, and stained with antibodies against Vps41. Arrows point to two neurons that either were transfected (top) or untransfected (bottom) with Vps41 shRNA. Scale bar = 10 μm. **m**, Quantitation of Vps41 signal intensity from (**l**). Box shows lower (Q1) and upper (Q3) quartile and median. Whiskers mark the 9 to 91 percentile range. Datapoints outside of this range are represented by individual dots. N_cells_ are shown inside boxplots. **n**, Representative axons from neurons co-transfected with PrP^PG14^-mCh and Vps41 siRNA, stained with antibodies against Vps41. Arrowheads indicate aggregate. Scale bars = 10 μm. **o**, Quantitation of Vps41 signal intensity from (**n**). Box shows lower (Q1) and upper (Q3) quartile and median. Whiskers mark the 9 to 91 percentile range. Datapoints outside of this range are represented by individual dots. N_axons_ are shown inside boxplots. **p**, Representative images of soma of neurons co-transfected with PrP^PG14^-mCh and shRNA scrambled or Vps41 siRNA. Scale bars = 10 μm. **q**, Quantitation of motile and stationary axonal PrP^PG14^-mCh vesicle densities in neurons co-transfected with scrambled or Vps41 shRNAs. N_vesicles_ are shown inside bars. All values in (**f, j, k, q**) are shown as mean ± SEM. Values in (**m, o**) are shown as Tukey’s box plot. *p<0.05, **p<0.01, ns = non-significant. Student’s t-test **(f, q**), Wilconxon rank-sum test (**m, o**).

**Supplementary Figure 7.**
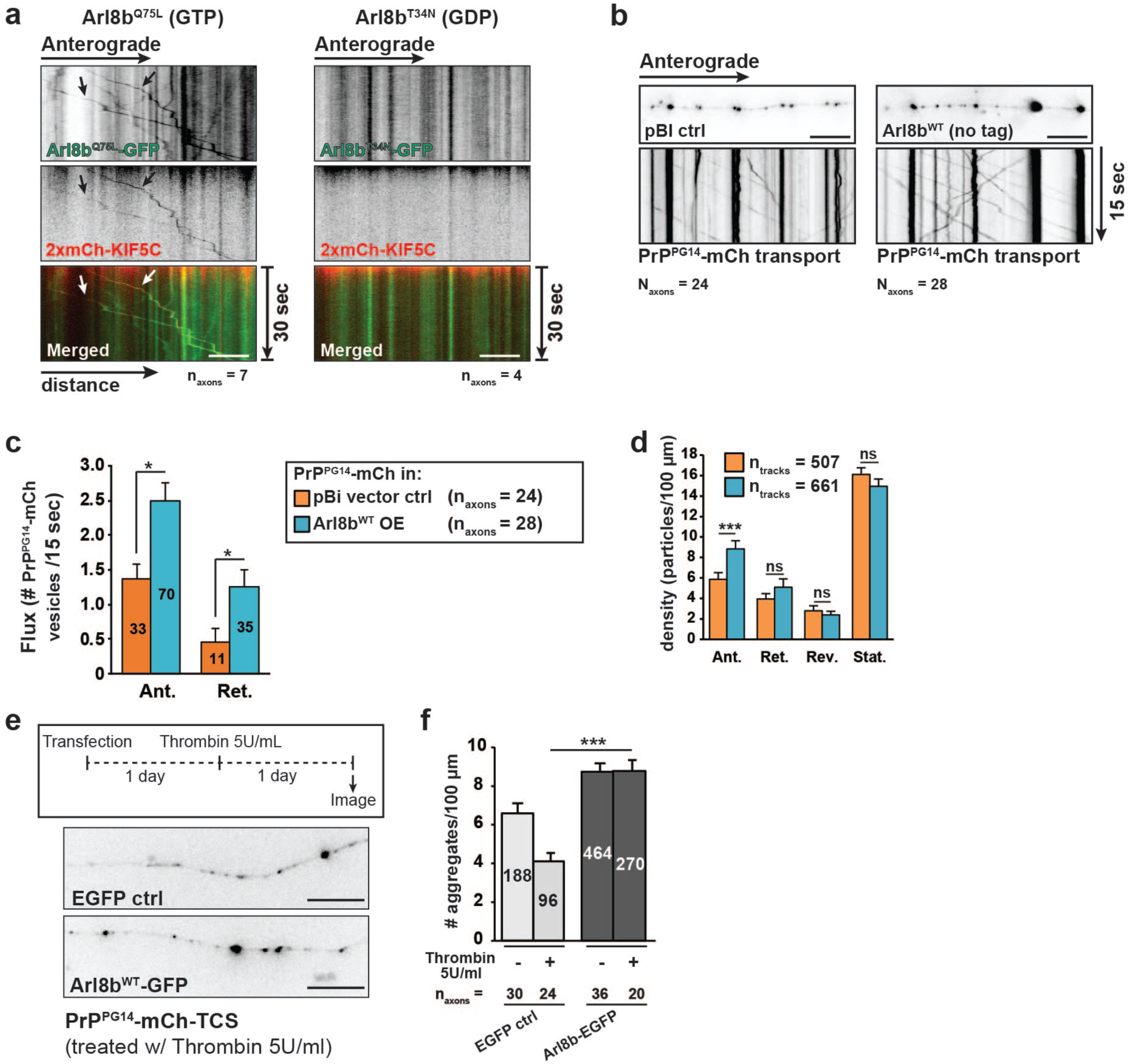
**Direct and indirect ARESTA act in parallel and converge on Arl8b. Related to Fig. 6. a**, Representative kymographs of axons co-expressing Arl8b^Q75L^-GFP (left) or Arl8b^T34N^-GDP (right) and 2xmCh-KIF5C. Arrows indicate cotransport. Scale bars = 10 μm. **b**, Representative first-frame images of time-lapse movies, and corresponding kymographs of axons from neurons expressing PrP^PG14^-mCh and transfected with empty pBi vector (control) or untagged Arl8b^WT^. Scale bars = 10 μm. **c**, Quantitation of PrP^PG14^-EGFP vesicle flux from (**b**). N_vesicles_ are shown inside bars. **d**, Quantitation of vesicle densities from (**b**). **e**, Top: Experimental outline of thrombin assay. Bottom: Representative images of axons of neurons co-expressing PrP^PG14^-mCh-TCS a. d soluble EGFP or Arl8b^WT^-GFP. Scale bar = 10 μm. **f**, Quantitation of PrP^PG14^-mCh aggregate densities from (**e**). N_aggregates_ are shown inside bars. Ant. = anterograde; Ret. = retrograde; Rev. = reversal; Stat. = stationary. Values in (**c, d, f**) are shown as mean ± SEM. *p<0.05, ***p<0.001, Kruskal-Wallis test.

**Supplementary Figure 8.**
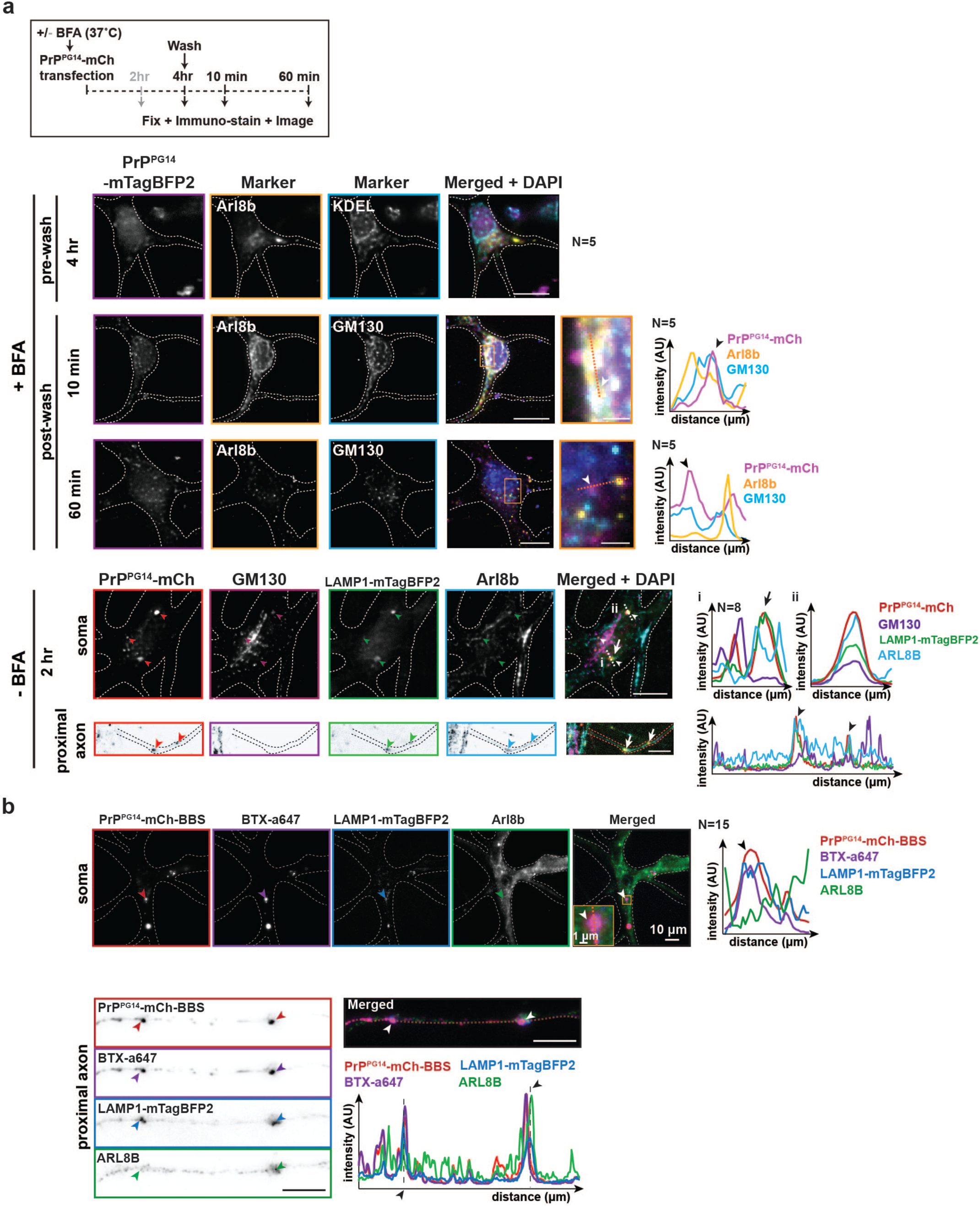
**Arl8b is a key determinant of PrP^PG14^ axonal entry. a**, Top panels : experimental outline of BFA assay. Bottom panels: representative images of soma and proximal axons of different neurons expressing PrP^PG14^-mTagBFP2 or PrP^PG14^-mCh + Lamp1-mTagBFP2 and stained with antibodies against ARL8B, ER (KDEL), Golgi (GM130), or with DAPI nuclear marker at indicated time points and conditions. Graphs to the right are of line scans of dotted lines inside insets. Arrowheads point to colocalization events, also shown in enlarged insets and in linescan intensity profiles. Scale bars in main panels = 10 μm. Scale bars in insets = 500 nm. **b**, Representative images PrP^PG14^-mCh-BBS internalization assay of the soma (top panels) and axons (bottom panels), of neurons co-transfected with PrP^PG14^-mCh-BBS and LAMP1-mTagBFP2, labeled with BTX-a647, and stained with an antibody against ARL8B. Arrowheads point to colocalization events, also shown in enlarged insets and in linescan intensity profiles. Scale bars in main panels = 10 μm. Scale bars in inset = 1 μm.

**Supplementary Figure 9.**
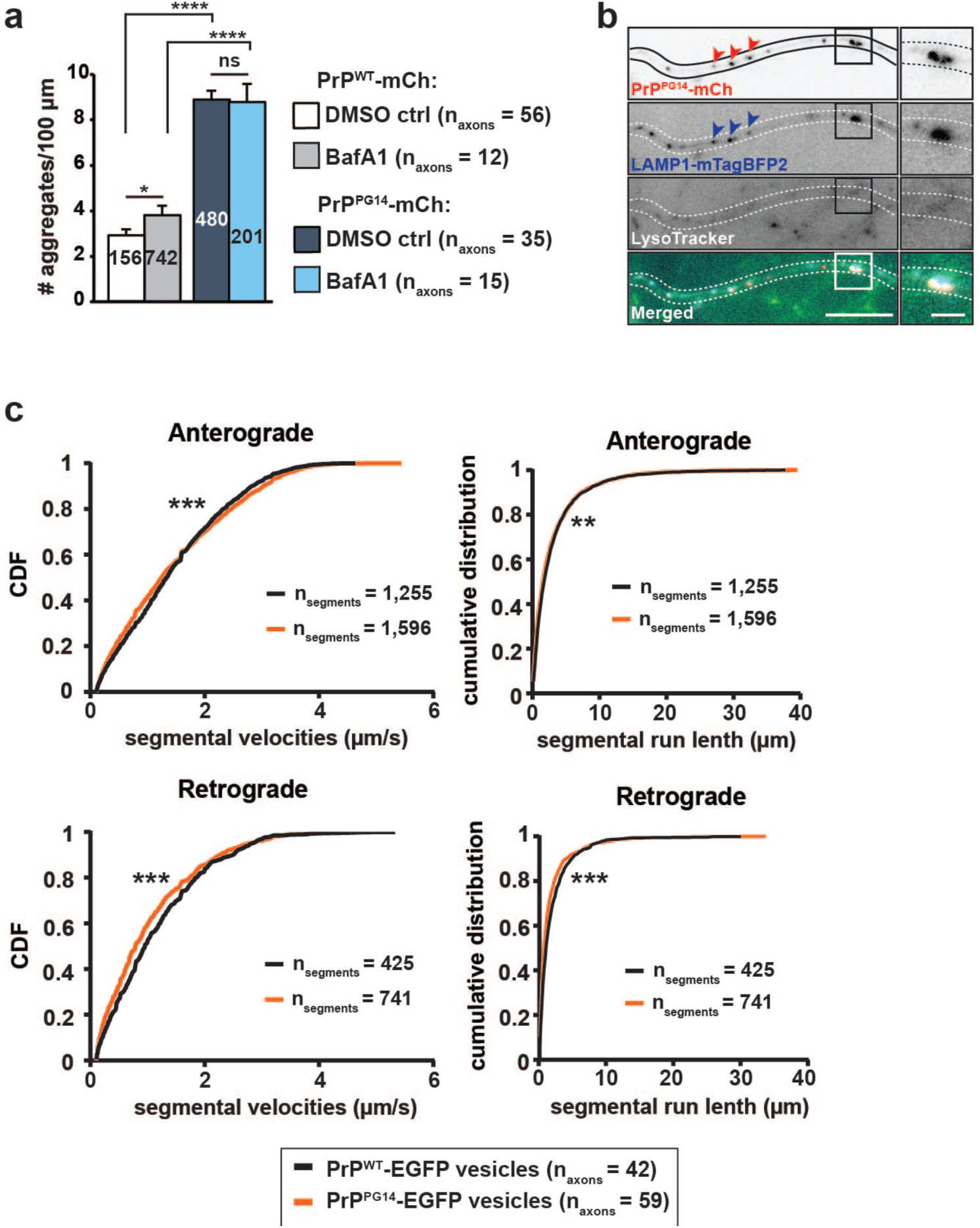
**Lysosomal degradation impairments and retrograde transport deficits in PrP^PG14^-expressing axons. Related to Figure 7. a**, Quantitation of axonal PrP^PG14^-mCh aggregate densities in neurons treated with BafA1. N_aggregates_ are shown inside bars. **b**, Representative images of axons of neurons co-expressing PrP^PG14^-mCh and LAMP1-mTagBFP2 and treated with LysoTracker. Arrowheads point to colocalization. Right panels are enlargements of boxed regions. N_axons_ = 21. Scale bar of main figure = 10 μm. Scale bar of inset = 2.5 μm. **c**, Cumulative distribution frequencies (CDFs) of anterograde (top) and retrograde (bottom) segmental velocities and segmental run length of PrP^WT^- and PrP^PG14^-EGFP vesicles. **p<0.01, ***p<0.001, Kolmogorov-Smirnov test.

**Supplementary Figure 10.**
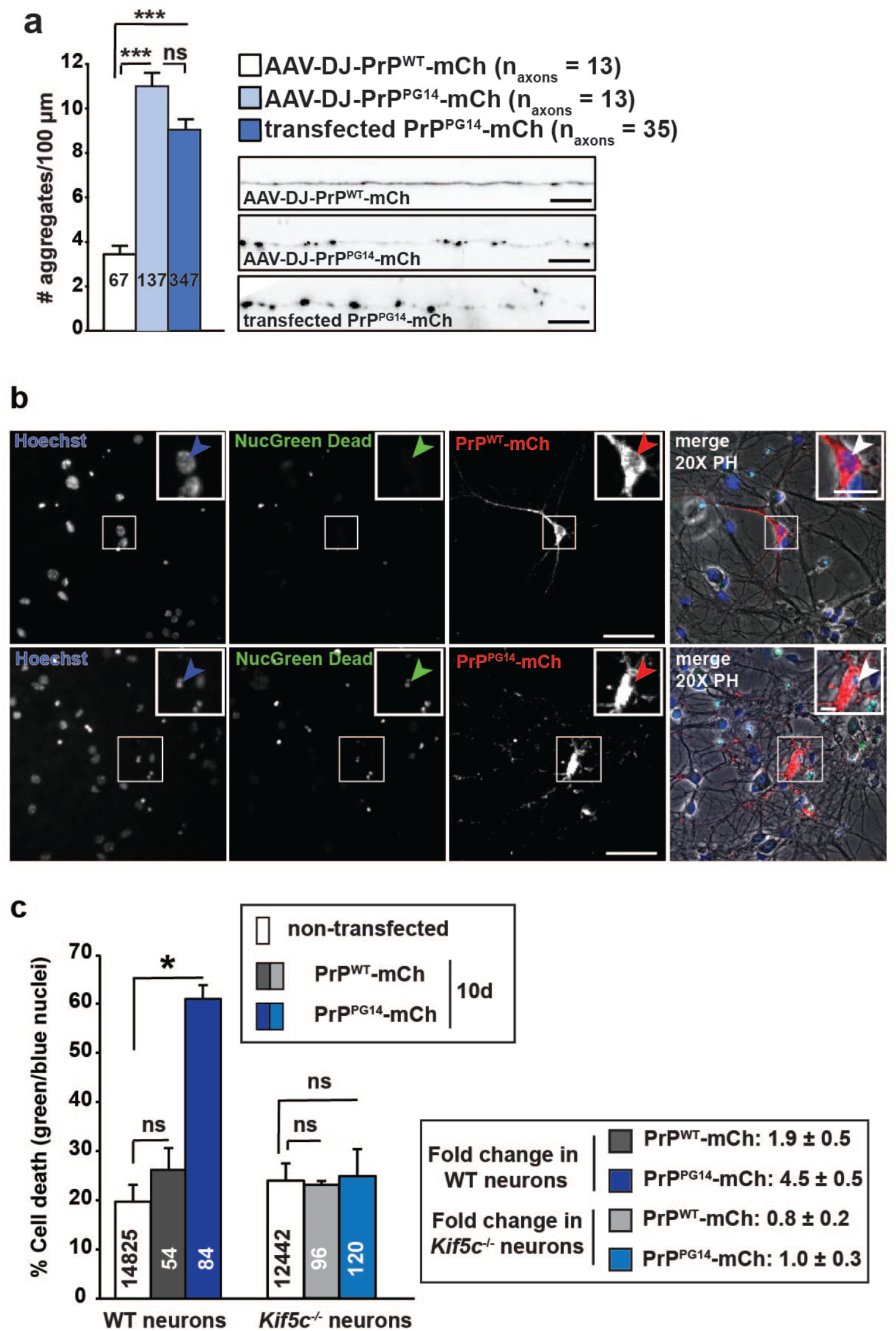
**Kinesin-1-dependent endoggresomes impair calcium influx and accelerate cell death of PrP^PG14^ neurons. Related to Figure 8. a**, Quantitation of aggregate densities (left) and representative images (right) of axons from neurons transduced with AAV-DJ-PrP^PG14^-mCh. N_aggregates_ are shown inside bars. Scale bar = 10 μm. **b**, Representative images of hippocampal neurons expressing PrP^WT^- or PrP^PG14^-mCh and stained with Hoechst and NucGreeen Dead. Arrowheads point to alive (top panels) and dead (bottom panels) neurons. Insets are enlarged in the respective top right corners of each panel. Scale bars on main figures = 100 μm. Scale bars on insets = 20 μm. **c**, Quantitation of percentage cell death from (**b**). N_cells_ are shown inside bars. Fold changes between indicated conditions are indicated on boxed panel (right). All values are shown as mean ± SEM. *p<0.05, ***p<0.001, ns = non-significant, Kruskal-Wallis test.

**Supplementary Figure 11.**
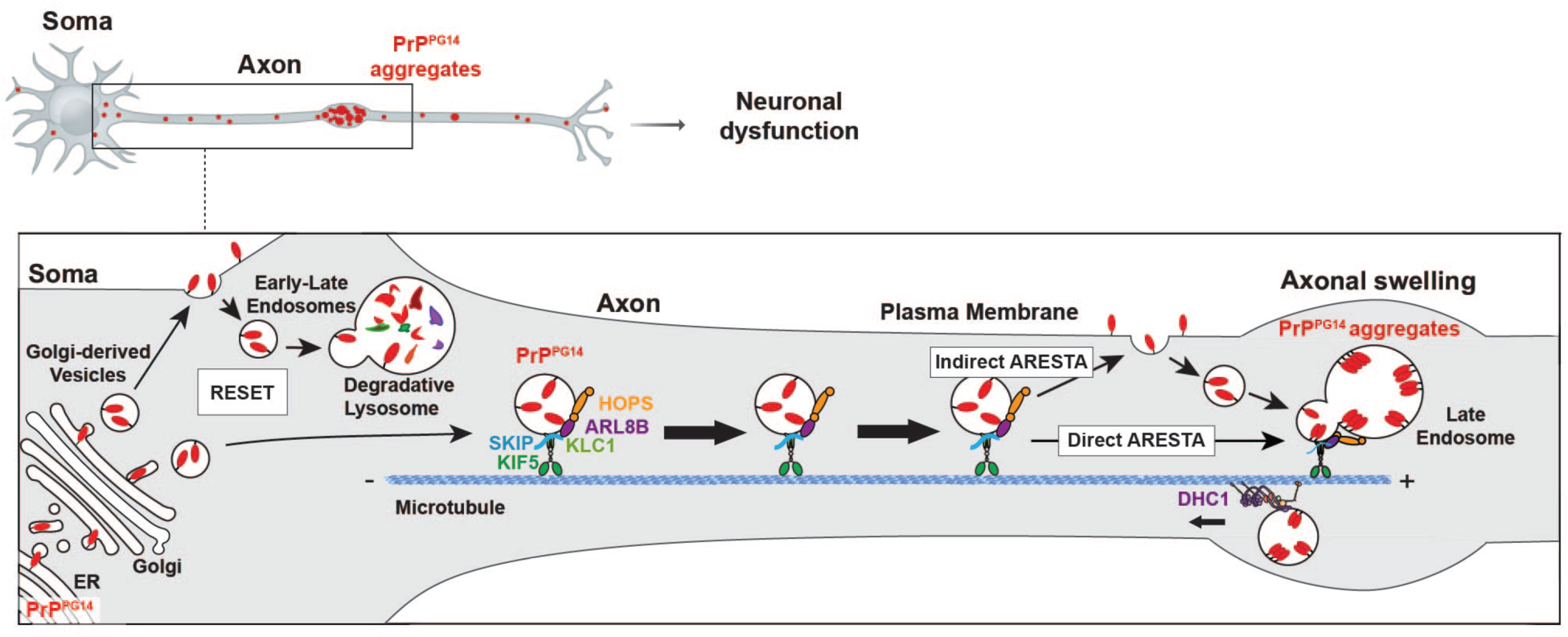
Neuronal endo-lysosomal trafficking RESET and ARESTA pathways that clear misfolded PrP or form endoggresomes. Model showing endosomal pathways governing mutant PrP degradation versus aggregation in neurons. RESET acts in the soma to degrade PrP^PG14^ via the cell surface. Arl8b/kinesin-1/SKIP/HOPS earmarks PrP^PG14^ vesicles as endosomes and drives their axonal entry and toward ARESTA-dependent aggregation. Aggregation occurs following indirect targeting of PrP^PG14^ to late endosomes via the axonal cell surface, via its direct homotypic fusion, or both. Dynein-mediated retrograde transport and axonal degradative capacity are impaired in axons thus promoting the maintenance of aggregates in axons.

## DATA AVAILABILITY

The datasets generated during and/or analyzed during the current study are available from the corresponding author on reasonable request.

